# A combinatorial code of neurexin-3 alternative splicing controls inhibitory synapses via a trans-synaptic dystroglycan signaling loop

**DOI:** 10.1101/2022.05.09.491206

**Authors:** Justin H. Trotter, Cosmos Yuqi Wang, Peng Zhou, Thomas C. Südhof

**Affiliations:** Dept. of Molecular and Cellular Physiology, Stanford University School of Medicine, Stanford, CA 94305, USA; Howard Hughes Medical Institute, Stanford University School of Medicine, Stanford, CA 94305, USA

## Abstract

Disrupted synaptic inhibition is implicated in most psychiatric disorders, yet the molecular mechanisms that shape and sustain inhibitory synapses are poorly understood. Here, we find that the function of a subset of inhibitory synapses in brain is controlled by binding of presynaptic Neurexin-3 to postsynaptic dystroglycan, adhesion molecules that are associated with cognitive impairments. We found that Neurexin-3 alternative splicing at two sites, SS2 and SS4, acts as an ‘and/or’ logic gate to regulate the release probability, but not the number, of inhibitory synapses in olfactory bulb and prefrontal cortex. The same SS2 and SS4 splice variants that enable inhibitory synapse function also allow binding of Neurexin-3 to dystroglycan. Inactivation of postsynaptic dystroglycan, in turn, produces a similar decrease in release probability as the presynaptic Neurexin-3 deletion. Furthermore, a minimal dystroglycan-binding construct of Neurexin-3 fully sustains inhibitory synaptic function, suggesting that trans-synaptic dystroglycan binding is not only necessary but also sufficient for inhibitory synapse function. Thus, Neurexin-3 enables a normal release probability at inhibitory synapses via a trans-synaptic feedback loop requiring binding of presynaptic Neurexin-3 to postsynaptic dystroglycan.

---

Synapses are sophisticated intercellular junctions controlled by trans-synaptic signaling that is mediated, at least in part, by interactions between pre- and postsynaptic adhesion molecules. Among synaptic adhesion molecules (SAMs), neurexins stand out because of their central role in shaping the properties of synapses^1–4^. Neurexins mediate a panoply of synaptic functions, ranging from mediating a normal neurotransmitter release probabiity^5–7^ to controlling presynaptic GABA_B_-receptors^8^, regulating postsynaptic glutamate and GABA_A_-receptors^9–12^, and enabling postsynaptic NMDA-receptor-dependent LTP^13^. Recent observations uncovered multitudinous interactions of neurexins with diverse ligands that likely mediate the functions of neurexins. Indeed, trans-synaptic ligands for some of these functions were identified, as shown for the neurexin-dependent control of glutamate receptors that is effected by Cbln1/2-GluD1/2 complexes in the subiculum^11^ or for neurexin-dependent LTP that requires *Nlgn1* in the hippocampal CA1 region^14^. Among the various neurexin functions, their most impactful role probably consists of their regulation of the presynaptic release probability^5–7^, but here no candidate ligands were identified, and it is unclear how neurexins determine the release probability of a synapse. Even the question whether presynaptic neurexins directly act cell-autonomously in the nerve terminal, or whether they operate indirectly via binding to a postsynaptic ligand that then signals back to the presynaptic release machinery remains unknown^4^.

Neurexins are transcribed from three genes in two principal forms, longer α-neurexins and shorter β-neurexins^15–17^. Neurexin mRNAs are extensively alternatively spliced at six canonical positions (SS1 to SS6)^18, 19^, whose use is highly regulated spatially and temporally^20, 21^. However, only one site of alternative splicing has been shown to be functionally relevant, splice site #4 (SS4), which regulates binding of key ligands to neurexins^1, 4, 22–25^ and controls the postsynaptic levels of AMPA-(AMPARs) and NMDA-receptors (NMDARs)^10^. In addition to SS4, only one other site of alternative splicing, SS2, has been found to modulate neurexin ligand interactions, in that SS2 regulates binding of α-neurexins (β-neurexins lack SS2) to dystroglycan, a postsynaptic adhesion molecule, and to neurexophilins, a family of secreted cysteine-rich proteins^26–30^. Strikingly, dystroglycan binds only to α-neurexins lacking an insert in either SS2 and/or in SS4, but not to α-neurexins with inserts in both SS2 and SS4^26, 30^. However, whether dystroglycan binding to α-neurexins is physiologically important is not known since in addition to neurexins, dystroglycan binds to a large number of potential ligands, such as agrin^31^, pikachurin^32^, and slit^33^. Indeed, circumstantial evidence from studies of CCK-positive synapses in the hippocampus suggested that dystroglycan-binding to neurexins is functionally insignificant^34^.

In brain, diverse types of inhibitory neurons form distinct types of inhibitory synapses^35, 36^, and dystroglycan is essential for the formation and/or function of a subset of these synapses^34, 37–40^. Mutations in genes that are part of the dystroglycan complex, such as dystrophin and LARGE (the enzyme that uniquely glycosylates dystroglycan), are often associated with cognitive impairments^41^. Thus, the function of dystroglycan at inhibitory synapses could account for cognitive symptoms in these patients, but the mechanism of dystroglycan’s function in inhibitory synapses is enigmatic. Similarly, neurexin gene mutations have been associated with cognitive impairments in human patients. In particular, neurexin-3 (*NRXN3*) mutations were found in a number of families with neuropsychiatric disorders^42, 43^, and *NRXN3* is a class 1 gene in the SFARI autism gene database (https://gene.sfari.org/database/gene-scoring/), but again how *NRXN3* mutations predispose to cognitive impairments is unclear.

The present study was motivated by the unexpected finding that neurexin-3 (*Nrxn3*) in mice is essential for the normal release probability of an inhibitory synapse in the olfactory bulb (OB), namely the granule cell→mitral cell (GC→MC) synapse that constitutes one half of granule cell-mitral cell dendrodendritic synapses^44^. Here we demonstrate that only Neurexin-3α (Nrxn3α), but not Neurexin-3β (Nrxn3β), supports GC→MC synaptic transmission, and that the function of Nrxn3α is controlled by a hierarchical splice code involving SS2 and SS4 of Nrxn3α, such that either SS2 or SS4 need to lack an insert. Strikingly, a minimal Nrxn3α construct containing only a single LNS2-domain without an insert in SS2 and that binds to dystroglycan fully supports GC→MC synaptic transmission; moreover, Nrxn3α performs a similar function in inhibitory synapses of the medial prefrontal cortex (mPFC) with the same dependence on LNS2-domain alternative splicing at SS2. Furthermore, both at GC→MC synapses and at inhibitory synapses of the mPFC, postsynaptic dystroglycan deletions produce a similar phenotype as the presynaptic *Nrxn3* deletions. Our data thus suggest that binding of presynaptic Nrxn3α to postsynaptic dystroglycan tunes the presynaptic release probability at inhibitory synapses. This function of the Nrxn3α/dystroglycan assembly is tightly regulated by Nrxn3α alternative splicing, which may explain-at least in part-why mutations in *Nrxn3* and in dystroglycan-associated proteins both induce cognitive impairments.

## RESULTS

### Nrxn3α, but not Nrxn3β, rescues inhibitory synapse function in *Nrxn3*-deficient OB neurons

Neurexin deletions at many synapses cause a decrease in release probability^5–7^. To explore the mechanisms involved, we first focused on one particular synapse, the inhibitory GC→MC synapse in the OB, where one particular neurexin, *Nrxn3*, is essential for a normal release probability^44^. We used rescue experiments to elucidate the Nrxn3 sequences required for the release probability at this synapses, and tested SS4 splice variants because nearly all known functions of neurexins depend on this site of alternative splicing.

We infected mixed neuron-glia cultures from the OB of *Nrxn3* cKO mice with lentiviruses expressing either inactive (ΔCre, as a control), active Cre-recombinase (Cre) alone, or Cre together with Nrxn3α^SS4+^ or Nrxn3α^SS4-^ (containing or lacking an insert in SS4, respectively), or with Nrxn3β^SS4+^ (containing an insert in SS4) (Fig. S1a). We then performed whole-cell patch-clamp recordings from larger mitral/tufted cells, which can be clearly distinguished from smaller inhibitory granule cells^45^, and used extracellular stimulations to evoke IPSCs. Since granule cells are by far the most abundant inhibitory neurons in the OB, these evoked IPSCs largely reflect GC→MC synaptic transmission in the cultured OB neurons, although they likely also contain a minor component derived from other inhibitory neurons in the cultures.

The *Nrxn3* deletion severely impaired (60-80% decrease) evoked IPSCs in cultured OB neurons (Fig. 1a, 1b), consistent with earlier results^44^. This impairment was rescued by expression of Nrxn3α^SS4+^ or Nrxn3α^SS4-^, which did not affect evoked IPSCs in control neurons (Fig. 1a, 1b, S1b). Nrxn3β^SS4+^, however, seemed to suppress evoked IPSCs both in control and in *Nrxn3*-deficient neurons, although in the initial experiments this effect was not statistically significant (Fig. 1a, 1b). Independent replication of this experiment confirmed that expression of Nrxn3β^SS4+^ in WT neurons decreased the evoked IPSC amplitude ∼50% (Fig. 1c, 1d). Measurements of evoked excitatory synaptic transmission, monitored as NMDAR-dependent evoked EPSCs, failed to detect any changes induced by the *Nrxn3* deletion or by the expression of Nrxn3α^SS4+^, Nrxn3α^SS4-^, or Nrxn3β^SS4+^, suggesting that the effect of the Nrxn3 deletion is specific for inhibitory synapses (Fig. 1e-1h, S1c).

**Figure 1:**
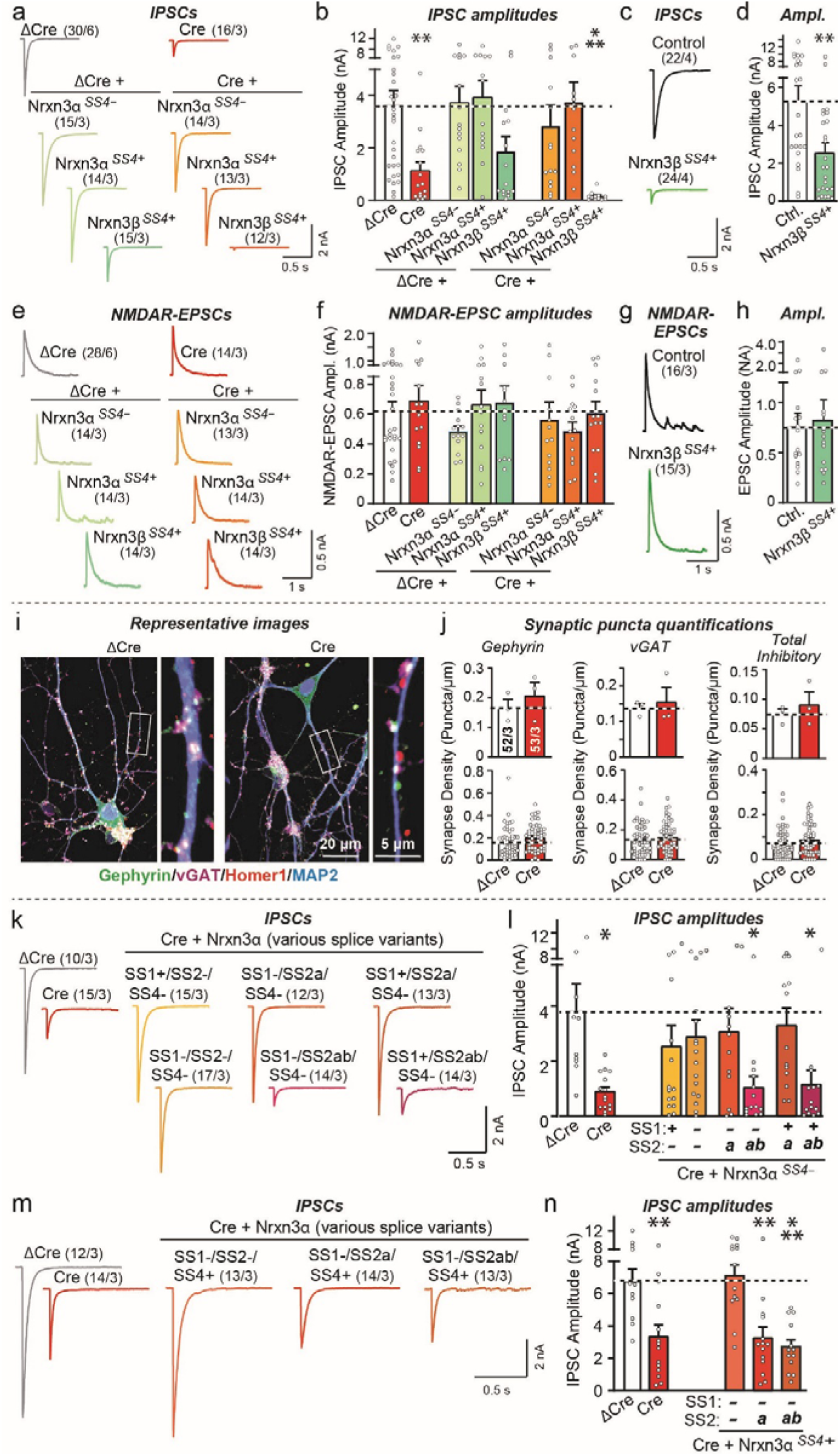
A hierarchical alternative splicing code of Nrxn3α at SS2 and SS4 regulates inhibitory synapse function in olfactory bulb neurons. All experiments were performed in dissociated olfactory bulb (OB) cultures from newborn *Nrxn3* cKO mice that were infected with lentiviruses expressing the ΔCre (control) or Cre without or with the indicated rescue constructs. **a** & **b**. Conditional deletion of *Nrxn3* severely impairs inhibitory synaptic transmission monitored via the evoked IPSC amplitudes; this decrease is rescued by Nrxn3α but not by Nrxn3β, whose overexpression instead appears to reduce IPSCs on its own (a, sample traces; b, summary graphs of IPSC amplitudes). **c** & **d**. Overexpression of Nrxn3β in wild-type OB neurons suppresses evoked IPSCs (c, sample traces; d, summary graphs of IPSC amplitudes). **e** & **f**. Conditional deletion of *Nrxn3* has no effect on evoked NMDAR-EPSCs (e, sample traces; f, summary graphs of EPSC amplitudes). **g** & **h**. Overexpression of Nrxn3β in wild-type OB neurons also does not impair evoked NMDAR-EPSCs (g, sample traces; h, summary graphs of IPSC amplitudes). **i** & **j**. Conditional deletion of *Nrxn3* does not change synapse numbers as analyzed by quantitative immunocytochemistry for a presynaptic (vGAT) for postsynaptic markers of inhibitory (gephyrin) and excitatory synapses (Homer1) (i, sample images; j, summary graphs of puncta densities plotted both as per experiment (top) or per region-of-interest (bottom), which boosts statistical significance independent of the number of experiments). For quantifications of the Homer1 signal, see Fig. S2. **k** & **l**. Nrxn3α lacking an insert in SS4 rescues impaired inhibitory synaptic transmission after conditional deletion of *Nrxn3* in OB neurons if SS2 contains no (Nrxn3α^SS2-^) or only a short insert (Nrxn3α^SS2a^), but not if SS2 contains a long insert (Nrxn3α^SS2ab^), whereas alternative splicing at SS1 has no effect (k, sample traces; l, summary graphs of IPSC amplitudes). **m** & **n**. Nrxn3α containing an insert in SS4 rescues impaired inhibitory synaptic transmission in *Nrxn3*-deficient OB neurons only if SS2 contains no insert (Nrxn3α^SS2-^), but not if SS2 carries either a short (Nrxn3α^SS2a^) or long insert (Nrxn3α^SS2ab^) (m, sample traces; n, summary graphs of IPSC amplitudes). Numerical data are means ± SEM; n’s (cells/experiments) are indicated above the sample traces (a-h, k-n) or in the summary graph bars (j) and apply to all graphs in an experimental series. Statistical analyses were performed with a one-way analysis of variance (ANOVA) with Dunnett’s multiple comparison test (b, f, l, and n) or a Welch’s *t* test (d, h, and j), with * = p<0.05, ** = p<0.01, and *** = p<0.001.

In agreement with the results on evoked IPSCs, deletion of *Nrxn3* greatly lowered (50-80% decrease depending on the experiment) the spontaneous mIPSC frequency but not the mIPSC amplitude in mitral/tufted cells in OB cultures (Fig. S1d-S1f). This decrease again was rescued by expression of Nrxn3α^SS4+^ or Nrxn3α^SS4-^ (Fig. S1g, S1h), whereas expression of Nrxn3β^SS4+^ suppressed the mIPSC frequency in *Nrxn3*-deficient neurons (Fig. S1g-S1j) and decreased the mIPSC frequency in wild-type neurons (Fig. S1k, S1l). The impairment in GC→MC synaptic transmission was not due to a decrease in synapse numbers induced by *Nrxn3* deletion because it had no effect on inhibitory synapse numbers (Fig. 1i-1j, S2).

Viewed together, these data suggest that at GC→MC synapses as modeled in cultured neurons, Nrxn3 is essential for synaptic transmission in a manner that can be rescued by Nrxn3α^SS4+^ or Nrxn3α^SS4-^ isoforms, but not by Nrxn3β^SS4+^ isoforms. Interestingly, RT-PCR measurements of the expression of neurexins and their SS4 splice variants revealed that the expression of Nrxn3α^SS4+^ is highly enriched in inhibitory OB neurons that are composed of more than 95% of granule cells as measured using RiboTag pulldowns of translating mRNAs in vGAT-positive neurons (Fig. S3). In contrast, Nrxn3β^SS4+^ and Nrxn3β^SS4-^ are nearly undetectable (Fig. S3). Thus, the dominant-negative action of Nrxn3β^SS4+^ likely does not physiologically operate, but may reflect a specific feature of reciprocal dendrodendritic synapses related to the need to avoid cis-interaction as observed for neurexins expressed by mitral cells^46^.

### A combinatorial splice code of SS2 and SS4 controls Nrxn3α function at GC→MC synapses in the OB as an ‘AND/OR’ logic gate

Since Nrxn3α rescued the *Nrxn3* KO phenotype independent of SS4 alternative splicing, we turned out attention to the only other alternatively spliced sequence of neurexins that is known to regulate ligand interactions, SS2. SS2 is present in the LNS2 domain of α-neurexins, which is lacking in β-neurexins^21^, and controls binding of neurexophilins and of dystroglycan to neurexins^26–30, 47–49^. SS2 is expressed in three variants, SS2-lacking an insert, SS2a containing an 8 amino-acid ‘a’ insert, and SS2ab containing an additional 7 amino-acid ‘b’ insert^21^.

RT-PCR measurements revealed that in most brain regions, *Nrxn1* and *Nrxn3* are predominantly expressed as SS2-variants, whereas approximately half of the *Nrxn2* mRNAs are present as SS2a variants (Fig. S4a-S4d). SS2ab variants are uniformly rare (<10%) for all neurexins and brain regions. In OB tissue and cultures, Nrxn3α is predominantly expressed as the Nrxn3α^SS2-^ variant, with Nrxn3α^SS2a^ accounting for ∼10% of transcripts, and Nrxn3α^SS2ab^ exhibiting levels of <5% of transcripts (Fig. S4a-S4d). Using RT-PCR with mRNAs isolated by RiboTag purification from inhibitory neurons (>95% of which are granule cells) and from mitral cells of the OB, we found that Nrxn3α is expressed in granule cells exclusively as the Nrxn3α^SS2-^ variant (>99%), whereas in mitral cells the Nrxn3α^SS2-^ and Nrxn3α^SS2a^ variants are produced almost equally (∼55% vs 45%; Fig. S4e, S4f). Thus, SS2 alternative splicing of Nrxn3 is highly regulated in the OB, but its pattern of alternative splicing is distinct from that of SS4 which is present as the SS4+ variant in the entire OB (Fig. S3).

Does alternative splicing at SS2 regulate the ability of Nrxn3α to rescue the *Nrxn3* KO phenotype in inhibitory synapses? To address this question, we probed the effects of both SS1 and SS2 alternative splicing, including SS1 alternative splicing in the analysis because SS1 is adjacent to SS2 and may have a peripheral effect on neurexophilin-binding to neurexins^29^, on GC→MC synaptic transmission in cultured neurons. Rescue experiments of *Nrxn3* KO neurons from the OB with different SS1, SS2 and SS4 variants of Nrxn3α uncovered a surprising finding: When SS4 lacked an insert, both Nrxn3α^SS2-^ and Nrxn3α^SS2a^ rescued, but Nrxn3α^SS2ab^ was inactive (Fig. 1k, 1l, S4g). When SS4 contained an insert, however, only Nrxn3α^SS2-^ but not Nrxn3α^SS2a^ or Nrxn3α^SS2ab^ was able to rescue (Fig. 1m, 1n, S4h). SS1 alternative splicing had no effect on rescue. Since *Nrxn3* is expressed in granule cells exclusively as the SS4+ variant (Fig. S3g), *Nrxn3* alternative splicing at SS2 thus controls *Nrxn3* function in GC→MC synapses of the OB, at least as monitored in cultured neurons. Overall, these data suggest that alternative splicing of Nrxn3α at SS2 and SS4 govern its function at inhibitory GC→MC synapses of the OB: SS2 or SS4 needs to lack an insert in order for Nrxn3α to sustain synaptic function, indicating that alternative splicing of Nrxn3α produces an ‘AND/OR’ logic gate like an electronic circuit for GC→MC synapses.

### Nrxn3α-LNS2^SS2-^ containing a single LNS-domain fully sustains GC→MC synapse function

We constructed a series of deletion constructs of Nrxn3α to determine which domains are required for inhibitory synapse function at GC→MC synapses (Fig. 2a). Rescue experiments with constructs that include various extracellular LNS- and EGF-domains revealed that a minimal Nrxn3α protein containing only a single LNS domain, the LNS2 domain in which SS2 is located, was sufficient to reverse the Nrxn3α KO phenotype (Fig. 2b-2g, S4i-S4k). In the minimal Nrxn3α-LNS2 constructs, the LNS2 domain is fused to the glycosylated stalk region of Nrxn3α that separates the last LNS domain from the membrane and that is followed by the transmembrane region and cytoplasmic tail (Fig. 2a). Importantly, the minimal Nrxn3α-LNS2 construct only rescued inhibitory synaptic transmission in *Nrxn3* KO neurons when SS2 in the LNS2-domain lacked an insert (Nrxn3α-LNS2^SS2-^). Even the introduction of the short ‘a’ insert (Nrxn3α-LNS2^SS2a^) completely abolished rescue (Fig. 2f, 2g). Thus, surprisingly, a single LNS domain of Nrxn3α is sufficient to maintain synaptic transmission, indicating that most domains comprising the Nrxn3α architecture are dispensable for its function within GC→MC synapses.

**Figure 2:**
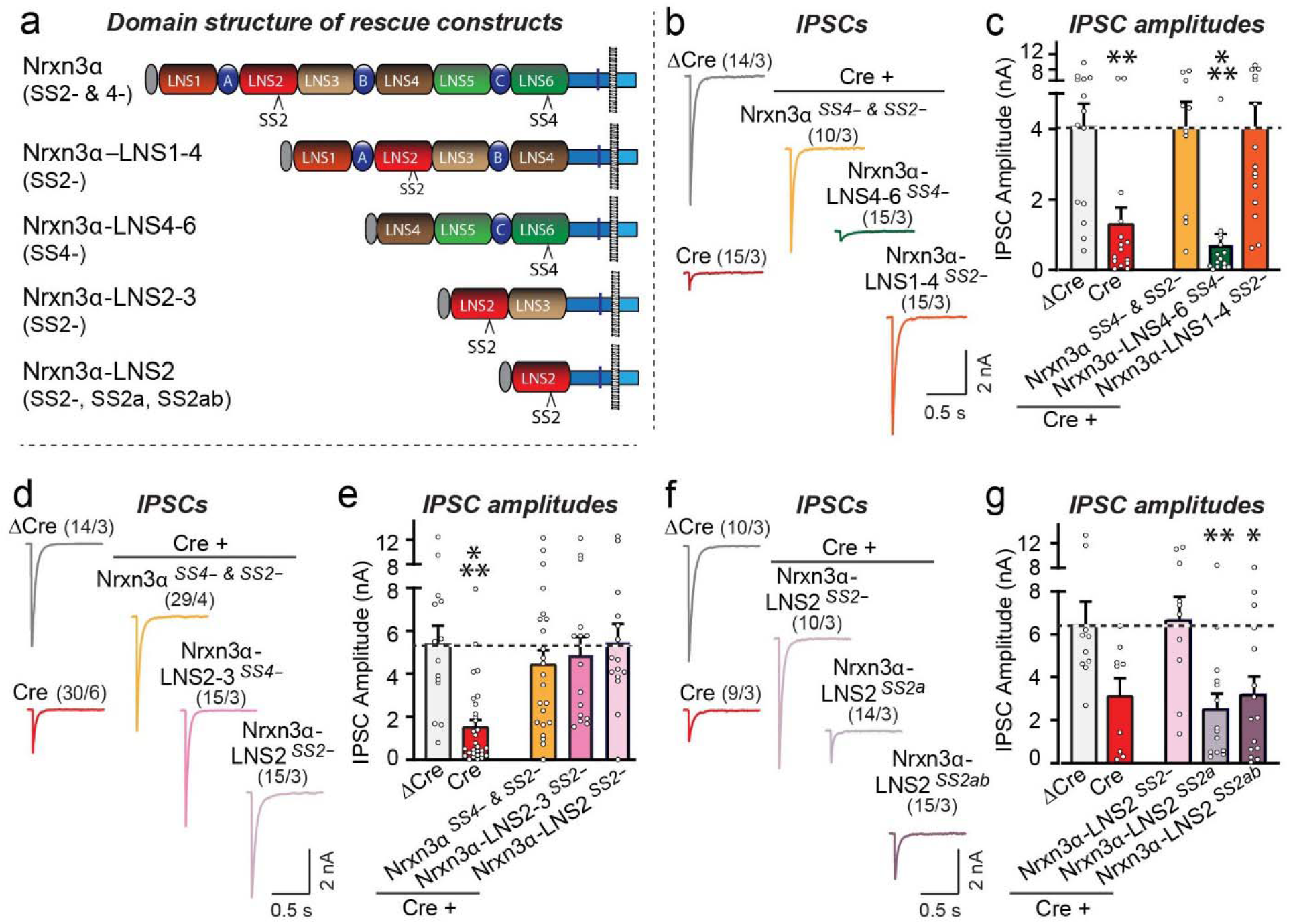
A minimal protein composed of the single LNS2-domain of Nrxn3α attached to its C-terminal transmembrane region and cytoplasmic sequence fully rescues inhibitory synaptic transmission in cultured *Nrxn3*-deficient OB neurons. All experiments were performed in dissociated OB cultures obtained from newborn *Nrxn3* cKO mice that were infected with lentiviruses to express ΔCre (control) or Cre without or with the indicated rescue proteins. **a**. Schematic of the Nrxn3α rescue constructs used. **b** & **c**. Nrxn3α lacking LNS5 and LNS6 domains fully rescues impaired inhibitory synaptic transmission in *Nrxn3*-deficient OB neurons (b, sample traces; c, summary graphs of IPSC amplitudes). **d** & **e**. A minimal Nrxn3α protein containing only LNS2 with the SS2-splice variant rescues impaired inhibitory synaptic transmission in *Nrxn3*-deficient OB neurons (d, sample traces; e, summary graphs of IPSC amplitudes). **f** & **g**. Rescue of the impaired inhibitory synaptic transmission in Nrxn3-deficient OB neurons by the minimal Nrxn3α protein containing only LNS2 is blocked if the SS2 site in LNS2 contains an insert (f, sample traces; g, summary graphs of IPSC amplitudes). Numerical data are means ± SEM; n’s (cells/experiments) are indicated above the sample traces and apply to all graphs in an experimental series. Statistical analyses were performed with a one-way analysis of variance (ANOVA) with Dunnett’s multiple comparison test (c, e, and g), with * = p<0.05, ** = p<0.01, and *** = p<0.001.

### Nrxn3α-LNS2^SS2-^ sustains GC→MC synaptic transmission in vivo without affecting synapse numbers

The finding that Nrxn3α-LNS2^SS2-^ may fully enable GC→MC synaptic transmission is surprising, raising the possibility that the preparation used – GC→MC synapses in cultured OB neurons – may be an epiphenomenon of dissociated cultures. Thus, we decided to examine the phenotype of the *Nrxn3* deletion and its rescue by Nrxn3α-LNS2^SS2-^ in GC→MC synapses in vivo.

We stereotactically infected the OB of *Nrxn3* cKO mice at P21 with AAVs encoding either ΔCre (as a control) or Cre (to delete *Nrxn3*), without or with Nrxn3α-LNS2^SS2-^ or Nrxn3α-LNS2^SS2a^ rescue constructs (Fig. 3a). ΔCre and Cre were expressed as tdTomato fusion proteins to visualize AAV-infected cells. Morphological studies of OB sections of infected mice at P35-42 showed that tdTomato was nearly ubiquitously expressed (Fig. 3b). The Nrxn3α-LNS2^SS2-^ and Nrxn3α-LNS2^SS2a^ constructs include an extracellular N-terminal HA-epitope tag, enabling us to visualize their expression by immunocytochemistry of OB sections (Fig. 3c). The two minimal Nrxn3α-LNS2 proteins were similarly distributed in the synaptic strata of the OB, confirming their expression and correct localization (Fig. 3c). Thus, this approach is well suited for probing *Nrxn3* function in the OB in vivo.

**Figure 3:**
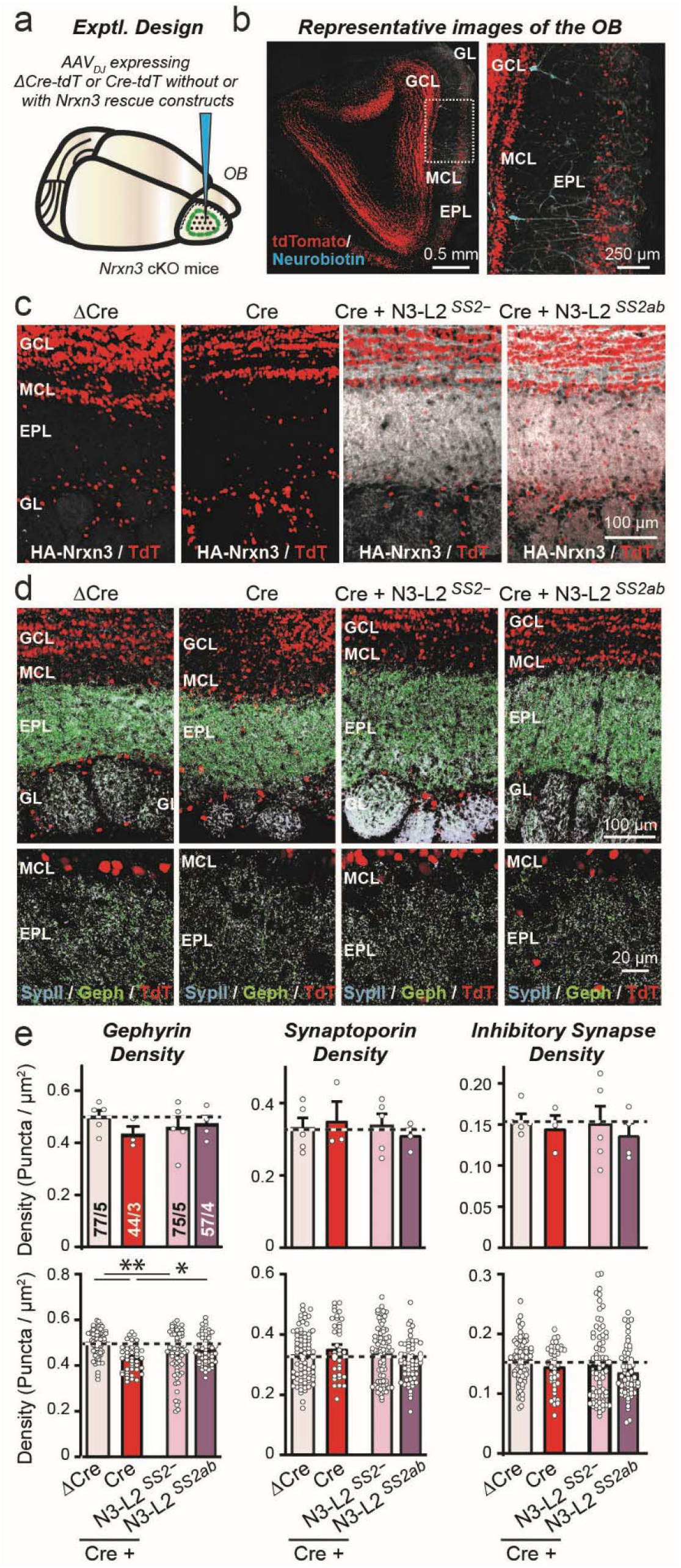
Conditional *Nrxn3* deletion in the OB *in vivo,* without or with rescue with minimal LNS2-domain *Nrxn3* constructs, has no effect on synapse density. **a** & **b**. Experimental design of in vivo *Nrxn3* deletions and rescues using stereotactic infections of the OB with AAVs expressing ΔCre (control), Cre, or Cre with additional AAVs expressing Nrxn3-LNS2 rescue constructs (a, schematic of stereotactic injections; b, representative fluorescence image of an OB section that was infected with AAVs expressing tdTomato fused to Cre, with subsequent patching of mitral cells that were filled with neurobiotin (blue)). Note that, as illustrated in b, the AAVs infect granule cells more efficiently than mitral cells. **c**. The minimal Nrxn3α-LNS2 rescue constructs are localized to synaptic layers in the OB after expression via AAVs, as visualized by immunocytochemistry for the N-terminal HA-epitope contained in the constructs (white, HA-Nrxn3α-LNS2 proteins; red, tdTomato). **d** & **e**. Conditional deletion of *Nrxn3* and rescue with minimal Nrxn3α-LNS2 constructs does not alter inhibitory synapse numbers *in vivo*. Sections from mice infected as shown in A were analyzed by quantitative immunocytochemistry for the presynaptic marker synaptoporin (a.k.a. synaptophysin-2; light blue) that is specific for granule cell→mitral cell synapses in the OB (Bergmann et al., 1993), and for the postsynaptic inhibitory synapse marker gephyrin (green) (d, sample images; e, summary graphs of puncta densities). Data are means ± SEM; n’s (cells/experiments) indicated in the summary graph bars apply to all graphs in an experimental series. Statistical analyses using one-way ANOVA with Dunnett’s multiple comparison test failed to uncover significant differences. Note that puncta densities in e are plotted both as analyzed per animal (top) and per region-of-interest (bottom); statistical significance is observed for gephyrin staining but not the other parameters when regions-of-interest are used as n’s because the pseudo-replicates in this analysis boost statistical significance independent of the actual number of experiments.

We next examined the effect of the *Nrxn3* KO and of the Nrxn3α-LNS2^SS2-^ and Nrxn3α-LNS2^SS2a^ rescue constructs in the OB on the density of GC→MC synapses. We stained OB sections for synaptoporin and for gephyrin, which are specific markers for GC→MC and for inhibitory synapses, respectively^50–52^, and analyzed the density of gephyrin- and synaptoporin-positive puncta as well as the density of puncta containing both signals (Fig. 3d, S5a). The results were analyzed using either the number of mice as the ‘n’ or the number of regions-of-interest (ROI’s) because the former is likely more correct, but the latter is the standard in the field despite the fact that the number of samples examined are not actually the number of replicates, but rather represent pseudo-replicates that boost statistical power without representing truly independent experimental observations. Using both statistical approaches, we detected no change in GC→MC synapse density except for a statistically significant 10% decrease using pseudo-replicate quantifications that was observed only for the gephyrin puncta density measurements (Fig. 3e, S5b). Viewed together, these results indicate that the *Nrxn3* KO in the OB does not cause a major change in synapse numbers.

Do in vivo deletions of *Nrxn3* cause an impairment in GC→MC synapse function that can be rescued by the minimal Nrxn3α-LNS2^SS2-^ construct, analogous to what we observed in cultured OB neurons? We addressed this critical question using whole-cell patch-clamp recordings from mitral cells in acute OB slices. No change in passive electrical properties of mitral cells were evident (Fig. S5c, S5d). Recordings of mIPSCs showed that the *Nrxn3* KO also robustly decreases (60%) the mIPSC frequency after an in vivo manipulation; this decrease was fully rescued by the Nrxn3α-LNS2^SS2-^ but not by the Nrxn3α-LNS2^SS2ab^ construct (Fig. 4a, 4b). No significant change in the mIPSC amplitude or kinetics were detected (Fig. 4c, S5e). A recent study found that Nrxn3 differentially regulates the formation and/or function of a subset of inhibitory synapses in the ventral subiculum of the hippocampus in a sex-dependent manner^53^. However, separate analyses of slices from male and female mice exhibited a similar decrease in mIPSC frequency (Fig. S5f), indicating that not all contributions of Nrxn3 to inhibitory synapse function are sex-dependent.

**Figure 4:**
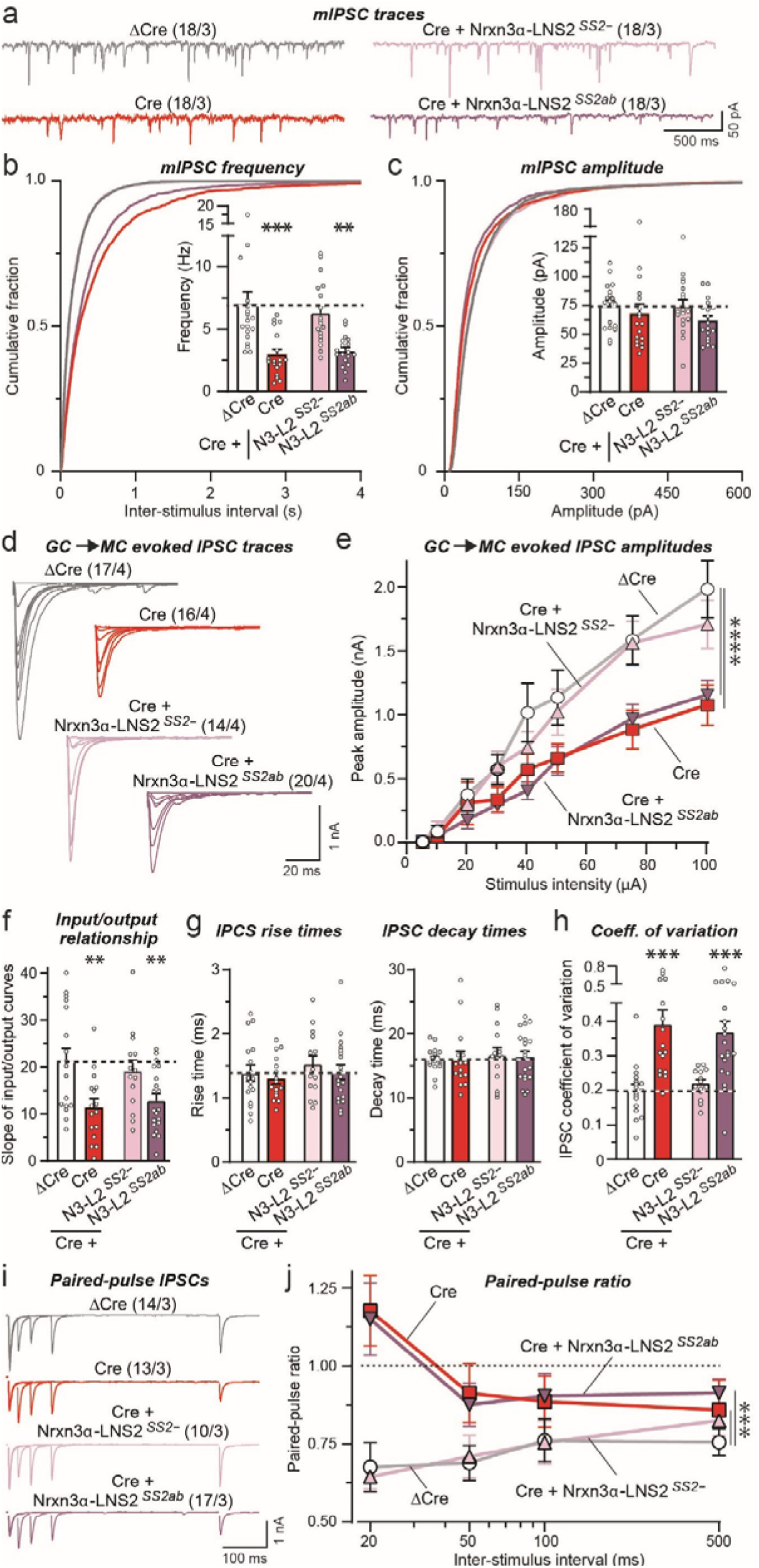
Conditional deletion of *Nrxn3* in the OB impairs GC→MC inhibitory synaptic transmission by lowering the release probability, which can be rescued by the minimal Nrxn3-LNS2 construct lacking an insert in SS2. All experiments were performed by patch-clamp recordings from mitral cells in acute slices sectioned from *Nrxn3* cKO mice whose OB was infected with AAVs (see Fig. 3a, S5c-S5e). **a**-**c**. The *Nrxn3* deletion decreases the mIPSC frequency in vivo; this decrease is rescued by the minimal Nrxn3α-LNS2 construct lacking an insert in SS2, but not by the same construct containing an insert in SS2 (a, representative mIPSC traces recorded in the presence of TTX; b, cumulative probability plots of the interevent interval and summary graph of the mIPSC frequency; c, cumulative probability plots and summary graph of the mIPSC amplitudes). **d**-**f**. The decrease in evoked IPSC amplitude induced by the *Nrxn3* deletion is rescued by the minimal Nrxn3α-LNS2 construct lacking an insert in SS2, but not by the same construct containing an insert in SS2 as documented by input/output curves to control for possible variations in stimulating electrode placement (d, representative IPSC traces; e, summary plot of input/output amplitude measurements; f, summary graph of the slope of the input/output curves). **g**. The *Nrxn3* deletion and expression of minimal Nrxn3α-LNS2 rescue constructs have no effect on the kinetics of evoked IPSCs (summary graphs of the IPSC rise (left) and decay times (right)). **h**. The *Nrxn3* deletion causes a massive increase in the coefficient of variation of IPSCs, suggesting a decrease in release probability, in a manner that can be rescued by the minimal Nrxn3α-LNS2 construct lacking an insert in SS2, but not by the same construct containing an insert in SS2. **i** & **j**. Consistent with a decreased release probability, the *Nrxn3* deletion induces a large increase in the paired-pulse ratio; this phenotype is also rescued by the minimal Nrxn3α-LNS2 construct lacking an insert in SS2, but not by the same construct containing an insert in SS2 (i, representative traces; j, summary plot of the paired-pulse ratio). Numerical data are means ± SEM; n’s (cells/experiments) are indicated above the sample traces and apply to all graphs in an experimental series. Statistical analyses were performed using two-way ANOVA in e and j and one-way ANOVA in b, c, and f-h with Dunnett’s multiple comparison test with regards to the ΔCre group, with * = p<0.05, ** = p<0.01, *** = p<0.001, and *** = p<0.0001.

To further probe synaptic transmission, we measured evoked IPSCs elicited by extracellular simulation. To control for possible effects caused by variations in the placement of the stimulating electrode, we analyzed evoked IPSCs as input/output curves (Fig. 4d). Again, the *Nrxn3* KO caused a large impairment (∼50% decrease) that was fully reversed by the Nrxn3α-LNS2^SS2-^ but not by the Nrxn3α-LNS2^SS2ab^ construct (Fig. 4d-4f). No change in IPSC kinetics was detectable (Fig. 4g). However, we found a large increase in the coefficient of variation of IPSCs that also was rescued by the Nrxn3α-LNS2^SS2-^ but not by the Nrxn3α-LNS2^SS2ab^ construct (Fig. 4h). This increase is indicative of a decrease in release probability^54^. To confirm a decrease in release probability, we measured the paired-pulse ratio of GC→MC IPSCs as a function of the interstimulus interval (Fig. 4i). The *Nrxn3* KO caused a massive increase in the paired-pulse ratio, again consistent with a decrease in release probability; this increase was also completely reversed by the Nrxn3α-LNS2^SS2-^ but not by the Nrxn3α-LNS2^SS2ab^ construct (Fig. 4j). Viewed together, these data show that the in vivo deletion of *Nrxn3* severely impairs neurotransmission at GC→MC synapses and that this impairment is due, at least in part, to a decrease in release probability. Moreover, consistent with our findings in cultured neurons, these data show that a minimal Nrxn3α construct containing only the LNS2 domain can rescue this impairment in a manner regulated by alternative splicing at SS2.

### *Nrxn3* deletion in the mPFC impairs inhibitory synaptic strength, which can be rescued via the minimal Nrxn3α-LNS2^SS2-^ construct

GC→MC synapses of the OB are a component of reciprocal dendrodendritic synapses that may differ from ‘standard’ inhibitory synapses in the CNS. The surprising finding that alternative splicing of Nrxn3α regulates this synapse via the activity of a single LNS domain could represent an exceptional mechanism that is specific to dendrodendritic synapses and not shared by other inhibitory synapses.

To ask whether the LNS2-dependent function of *Nrxn3* at GC→MC synapses of the OB may apply to other types of inhibitory synapses, we initially analyzed cultured cortical neurons. The *Nrxn3* KO nearly halved the amplitude and synaptic charge transfer of evoked IPSCs in cortical neurons (Fig. S6a, S6b). This decrease was rescued both by full-length Nrxn3α lacking inserts in SS2 and SS4, and by the minimal Nrxn3α-LNS2^SS2-^ construct (Fig. S6a, S6b).

Next, we deleted *Nrxn3* from mPFC neurons in vivo using stereotactic injections of AAVs expressing ΔCre (as a control) or Cre, with or without co-expression of the active Nrxn3α-LNS2^SS2-^ construct similar to the in vivo OB experiments (Fig. 5a). 2-3 weeks after infection, we sectioned acute slices from the mice and patched Layer 5/6 neurons for electrophysiological recordings (Fig. 5b). Measurements of spontaneous mIPSCs uncovered a robust decrease in mIPSC frequency (∼25%) induced by the *Nrxn3* KO (Fig. 5d), suggesting that a subset of the heterogeneous inhibitory synaptic inputs on Layer 5/6 neurons may have been impaired by the *Nrxn3* KO similar to GC→MC synapses. Expression of Nrxn3α-LNS2^SS2-^ fully rescued the decrease in mIPSC frequency in *Nrxn3* KO synapses. Moreover, we observed a small decrease in the mIPSC amplitude induced by the *Nrxn3* KO that was not rescued by the Nrxn3α-LNS2^SS2-^ construct (Fig. 5e), suggesting a different modest role for *Nrxn3* in postsynaptic GABA_A_R function in the mPFC. No changes in the passive electrical properties of the pyramidal mPFC neurons or in the mIPSC kinetics were detected (Fig. S6c-S6e).

**Figure 5:**
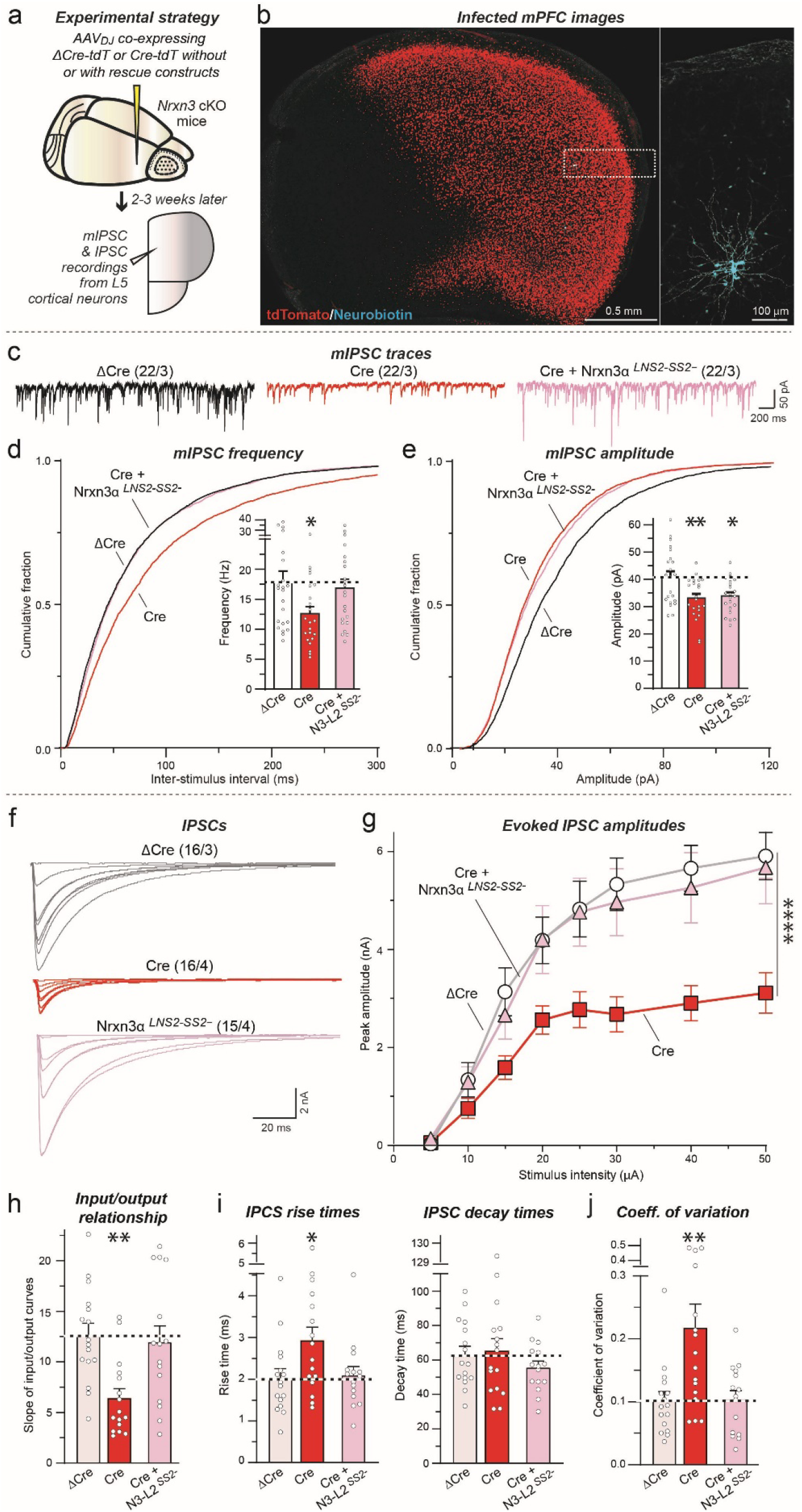
Deletion of *Nrxn3* in the mPFC impairs inhibitory synaptic function in a manner dependent on alternative splicing at SS2. **a.** Design of in vivo *Nrxn3* deletions using stereotactic infections of the mPFC with AAVs expressing ΔCre (control), Cre, or Cre with the minimal Nrxn3α-LNS2 rescue constructs. **b.** Representative fluorescence image of an mPFC section from a mouse infected with AAVs expressing Cre-tdTomato (red) in which a layer 5 pyramidal neuron was patched and filled with neurobiotin (expanded right image; blue). **c**-**e**. The *Nrxn3* deletion decreases the mIPSC frequency *in vivo*; this decrease is rescued by the minimal Nrxn3α-LNS2 construct lacking an insert in SS2 (c, representative mIPSC traces recorded in the presence of TTX; d, cumulative probability plots of the interevent interval and summary graph of the mIPSC frequency; e, cumulative probability plots and summary graph of the mIPSC amplitudes). **f**-**h**. The *Nrxn3* deletion greatly decreases the amplitude of IPSCs evoked by extracellular stimulation in layer 5 and recorded from pyramidal neurons in layer 5; again, this phenotype is rescued by the minimal Nrxn3α-LNS2 construct lacking an insert in SS2, as documented by input/output curves to control for possible variations in stimulating electrode placement (f, representative IPSC traces; g, summary plot of input/output amplitude measurements; h, summary graph of the slope of the input/output curves). Note that the input/output curves are not linear above 20 μA stimuli, suggesting a saturable number of axons from inhibitory neurons that form synapses on layer 5 neurons. **i**. The *Nrxn3* deletion increases the rise time of evoked IPSCs (left) in a manner that can be rescued by the minimal Nrxn3α-LNS2 rescue constructs. In contrast, *Nrxn3* deletion and rescue construct expression have no effect on decay times (right) of evoked IPSCs. **j**. The *Nrxn3* deletion causes a substantial elevation in the coefficient of variation of IPSCs, suggesting a decrease in release probability, in a manner that can be rescued by the minimal Nrxn3α-LNS2 construct lacking an insert in SS2. Numerical data are means ± SEM; n’s (cells/experiments) are indicated above the sample traces and apply to all graphs in an experimental series. Statistical analyses were performed using two-way ANOVA in g and one-way ANOVA in d, e, h-j with Dunnett’s multiple comparison test with regards to the ΔCre group, with * = p<0.05, ** = p<0.01, and *** = p<0.0001.

Next, we monitored evoked IPSCs, again using input/output measurements to control for possible variations in the placement of the stimulating electrode, even though -as always-all experiments were conducted ‘blindly’ (Fig. 5f). The *Nrxn3* deletion greatly suppressed the synaptic strength of evoked IPSCs (∼50% decrease), which could be fully rescued by the Nrxn3α-LNS2^SS2-^ construct (Fig. 5g, 5h). In addition, the *Nrxn3* KO caused a slowing of the IPSC rise but not decay times, again with full rescue by Nrxn3α-LNS2^SS2-^ (Fig. 5i). Finally, the *Nrxn3* KO induced a large increase (∼120%) in the coefficient of variation of evoked IPSCs in mPFC synapses similar to OB GC→MC synapses, and this phenotype was also completely reversed by Nrxn3α-LNS2^SS2-^ (Fig. 5j). Together these data indicate that Nrxn3α performs a similar function in a subset of inhibitory synapses in the mPFC as in GC→MC synapses in the OB, namely an essential role in sustaining the normal release probability at these synapses such that the *Nrxn3* deletion ablates nearly half of the inhibitory synaptic strength in a manner that can be rescued by the Nrxn3α-LNS2^SS2-^ construct.

### CRISPRi-mediated inhibition of dystroglycan expression phenocopies the *Nrxn3* KO at GC→MC synapses in cultured OB neurons

The requirement and sufficiency of the minimal Nrxn3α-LNS2^SS2-^ construct that only contains a single extracellular interaction domain (LNS2) with a specific splice variant (SS2-) for synaptic transmission at a subset of inhibitory synapses suggests Nrxn3α functions by binding to a trans-synaptic ligand. At present, only one ligand is known to specifically bind to the LNS2 domain of neurexins lacking an insert in SS2: dystroglycan^26, 30^. Neurexophilin also binds to the LNS2 domain, but its binding is enhanced instead of impeded by an insert in SS2^27–29^. Notably, dystroglycan also binds to the the LNS6 domain of α-neurexins when the LNS6 domain lacks an insert in SS4, accounting for the finding that full-length Nrxn3α is still functional at GC→MC synapses when it contains a partial insert in SS2 (i.e., SS2a^21^) as long as SS4 lacks an insert (Fig. 1)^26^.

To explore the possibility that dystroglycan may be the postsynaptic ligand for presynaptic Nrxn3α at GC→MC synapses that is required for sustaining their release probability, we selected a guide RNA (gRNA) that enables potent CRISPR interference (CRISPRi)-mediated inhibition of dystroglycan expression in cultured OB neurons (∼65% decrease in dystroglycan mRNA levels; Fig. 6a). Electrophysiological recordings revealed that the CRISPRi-mediated partial inhibition of dystroglycan expression induced a corresponding decrease (∼60%) in the amplitude of evoked IPSCs (Fig. 6b, 6c).

**Figure 6:**
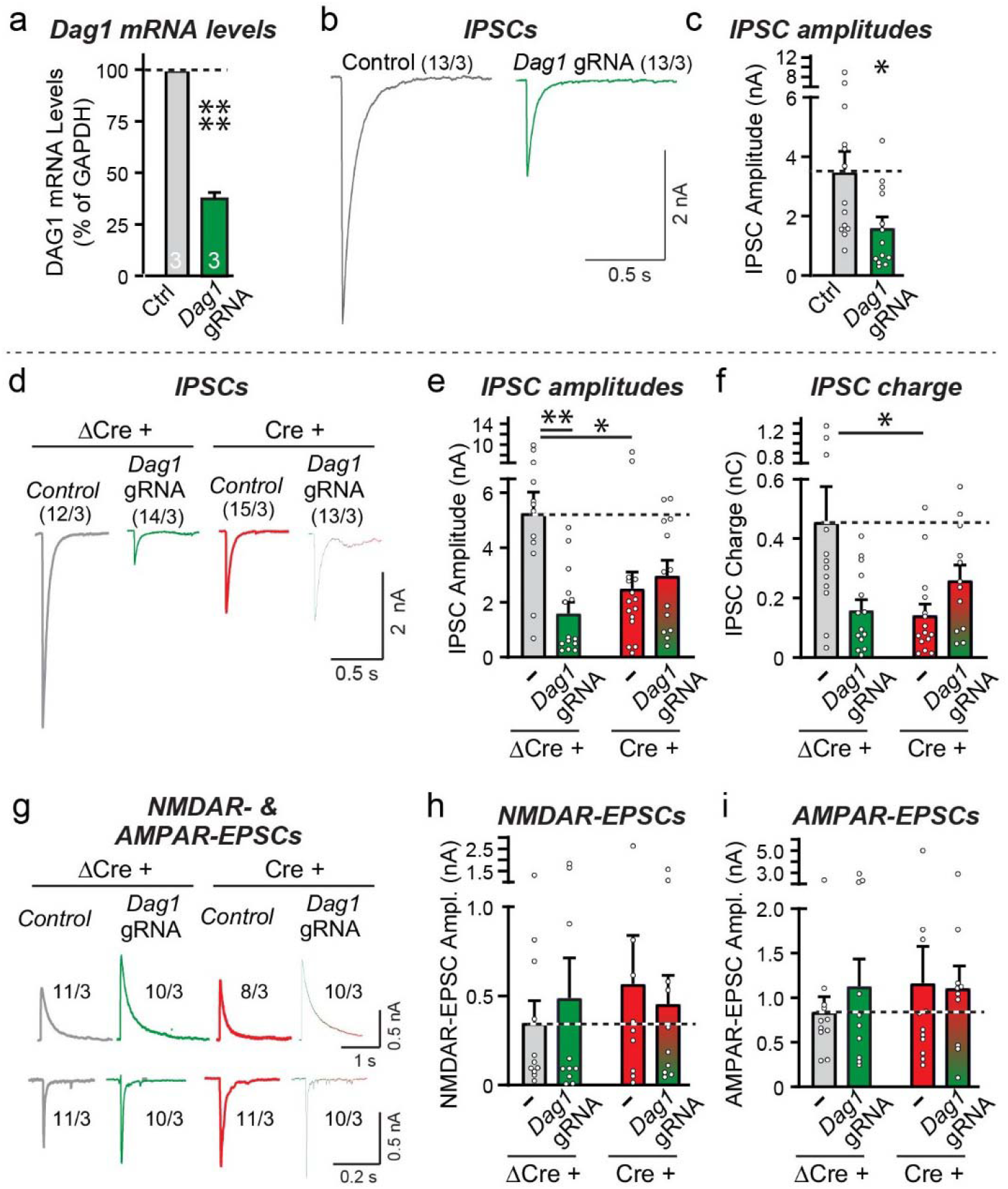
CRISPRi-mediated inhibition of dystroglycan expression in dissociated OB neurons selectively suppresses inhibitory synaptic transmission, with the *Nrxn3* and dystroglycan manipulations each occluding the other’s phenotype. **a-c**. CRISPR interference (CRISPRi)-mediated inhibition of dystroglycan (*Dag1*) expression partly suppresses the levels of dystroglycan mRNAs and significantly decreases the amplitude of evoked IPSCs (a, qRT-PCR measurements of Dag1 mRNA levels; b, sample traces; c, summary graph of IPSC amplitudes). **d**-**f**. Combined inhibition of dystroglycan (*Dag1*) expression and deletion of *Nrxn3* does not lower the evoked IPSC amplitude more severely than the single inhibition of expression or deletion of either dystroglycan or *Nrxn3*, respectively (d, sample traces; e & f, summary graphs of the IPSC amplitudes (e) and charge transfer (f)). IPSCs evoked by extracellular stimulation were recorded from mitral/tufted cells in dissociated culture obtained from *Nrxn3* cKO mice that were infected with lentiviruses expressing either ΔCre and/or the *Dag1* CRISPRi components. **g**-**i**. The single or double interferences with dystroglycan (*Dag1*) and of *Nrxn3* expression have no effect on evoked NMDAR- and AMPAR-EPSC amplitudes (g, sample traces; h & i, summary graphs of the evoked NMDAR-EPSC amplitudes (h) and AMPAR-EPSC amplitudes (i)). Experiments were performed as in d-f. Numerical data are means ± SEM; n’s (cells/experiments) are indicated above the sample traces and apply to all graphs in an experimental series. Statistical analyses were performed with a one-way analysis of variance (ANOVA) with Dunnett’s multiple comparison test (e, f, h, and i) or a Welch’s *t* test (a and c), with * = p<0.05, ** = p<0.01, *** = p<0.001, and **** = p<0.0001.

The suppression of IPSCs by the inhibition of dystroglycan expression in cultured OB neurons (Fig. 6b, 6c) is similar to that observed for the *Nrxn3* KO (Fig. 1a, 1b). To test whether these two adhesion molecules operate in the same pathway, we compared the phenotypes of single and double dystroglycan and *Nrxn3* KOs by combining the conditional deletion of *Nrxn3* with the CRISPRi-mediated inhibition of dystroglycan expression in cultured OB neurons from *Nrxn3* cKO mice. We infected the neurons with lentiviruses expressing either ΔCre (control) or Cre (to delete *Nrxn3*) and/or dCAS9-KRAB and the dystroglycan gRNA, and measured evoked IPSCs and NMDAR- and AMPAR-mediated EPSCs. The dystroglycan and *Nrxn3* deletions individually and together induced a 60-70% decrease in IPSC amplitudes and charge transfer, with no aggravation of the phenotype by the combined deletion compared to the individual deletions (Fig. 6d-6f). None of the deletions, individually or combined, had a significant effect on NMDAR- or AMPAR-EPSCs (Fig. 6g-6i). These data suggest that *Nrxn3* and dystroglycan act in the same pathway, consistent with the notion that they function by binding to each other.

### Postsynaptic dystroglycan deletion in vivo recapitulates the *Nrxn3* KO phenotype in the OB and mPFC

To validate the results obtained with the inhibition of dystroglycan expression in cultured OB neurons, we next investigated the effect of CRISPR-mediated deletion of dystroglycan in vivo both in the OB and the mPFC (Fig. 7, 8). We used CRISPR-mediated deletions instead of CRISPRi in these experiments because the components needed for CRISPRi could not be encoded by a single AAV. In the first set of experiments, we infected the OB of CAS9-expressing mice with AAVs encoding the dystroglycan gRNA or a control gRNA and tdTomato, and examined the efficiency of the dystroglycan deletion and the effect of the deletion on the inhibitory synapse density (Fig. 7a-7d, S7a-S7c). As assessed by immunocytochemistry for dystroglycan, the CRISPR-mediated dystroglycan deletion was efficient with a ∼60% decline in total dystroglycan signal (Fig. S7a, S7b). Contrasting prior reports that dystroglycan regulates the number of CCK+ inhibitory synapses in the hippocampus^34, 40^, quantifications of the density of inhibitory synapses visualized via immunocytochemistry for gephyrin failed to detect any change in synapse numbers or size in the external plexiform layer, the area that contains GC→MC synapses (Fig. 7c, 7d, S7c).

**Figure 7:**
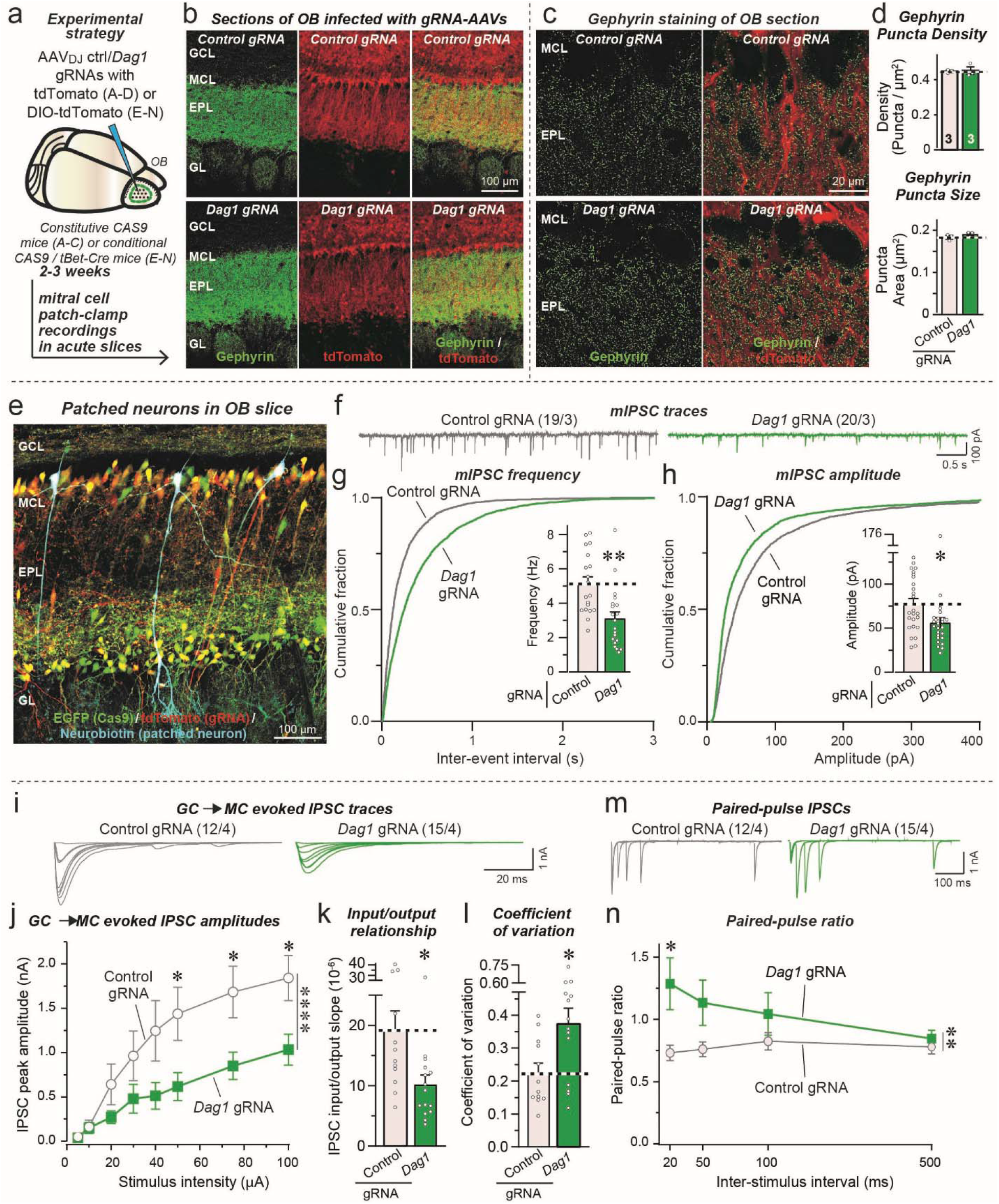
*In vivo* CRISPR-mediated deletion of dystroglycan (*Dag1*) via direct infection of the OB with AAVs expressing *Dag1* gRNAs decreases inhibitory GC→MC synaptic transmission by suppressing the release probability, but does not alter inhibitory synapse numbers. **a.** Experimental design. The OB of constitutive CAS9-expressing mice was infected stereotactically with AAVs encoding control or *Dag1* gRNAs together with tdTomato at P15-18, and mice were analyzed 2-3 weeks later. **b.** Representative fluorescence images of OB sections stained for gephyrin as an inhibitory synapse marker (green) and tdTomato expressed by the AAVs (red). **c** & **d**. Dystroglycan (*Dag1*) deletion in the OB *in vivo* does not change the density or size of gephyrin-positive synaptic puncta. Sections from mice infected as shown in a and b were analyzed by quantitative immunocytochemistry for the postsynaptic inhibitory synapse marker gephyrin (green) (c, sample images; d, summary graphs of puncta densities (top) and size (bottom)). Puncta densities and sizes are plotted as analyzed per animal, not per region-of-interest. **e**. Representative fluorescence image of a mitral cell filled with neurobiotin (blue) via the patch pipette in an OB section from a mouse with CRISPR-induced dystroglycan deletion in the OB (tdTomato expressed by the AAVs is shown in red, and EGFP expressed via the CAS9 knockin in green). **f**-**h**. Dystroglycan deletion in the OB *in vivo* decreases the mIPSC frequency monitored in mitral cells (f, representative mIPSC traces recorded in the presence of TTX; g, cumulative probability plots of the interevent interval and summary graph of the mIPSC frequency; h, cumulative probability plots and summary graph of the mIPSC amplitudes). **i**-**k**. Dystroglycan deletion in the OB *in vivo* suppresses inhibitory GC→MC synaptic transmission evoked by extracellular stimulation, as documented by input/output curves to control for variations in stimulating electrode placement (i, representative IPSC traces; j, summary plot of input/output amplitude measurements; k, summary graph of the slope of the input/output curves). **l**. Dystroglycan deletion in the OB *in vivo* causes a massive increase in the coefficient of variation of evoked IPSCs at GC→MC synapses, suggesting a decrease in release probability. **m** & **n**. Consistent with a decreased release probability, the *Dag1* deletion induces a large increase in the paired-pulse ratio (m, representative traces; n, summary plot of the paired-pulse ratio). Numerical data are means ± SEM; n’s (cells/experiments) are indicated in the summary graph bars (d) or above the sample traces (f, i and m) and apply to all graphs in an experimental series. Statistical analyses were performed using Student’s t-test in d, g, h, k, l and two-way ANOVA in j & n, with * = p<0.05, ** = p<0.01, *** = p<0.001, and **** = p<0.0001.

**Figure 8:**
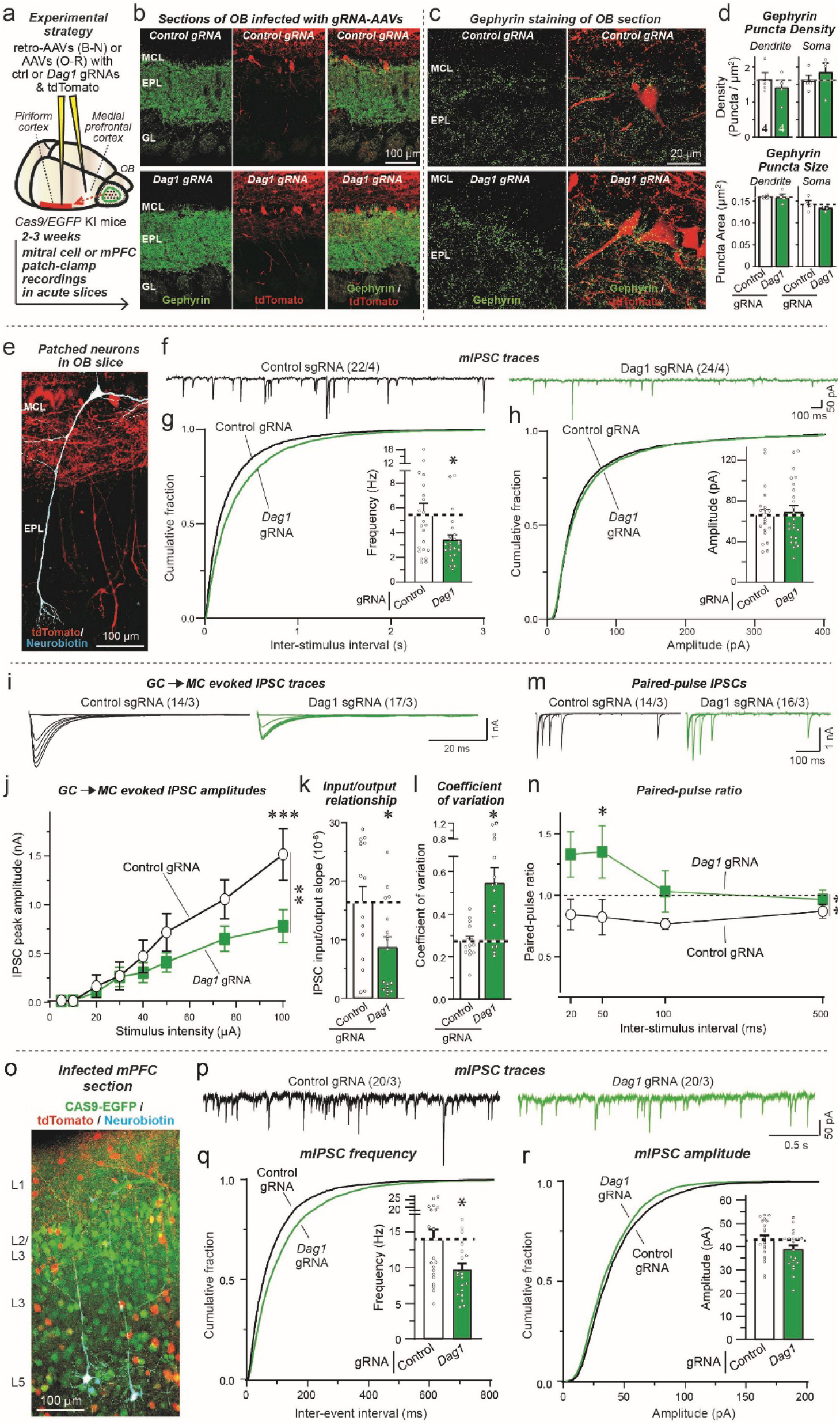
CRISPR-mediated *in vivo* deletion of dystroglycan (*Dag1*) via gRNA expression by retro-AAVs injected into the piriform cortex of CAS9 mice decreases inhibitory GC→MC synaptic transmission by suppressing the release probability without altering inhibitory synapse numbers, and a similar dystroglycan deletion in the mPFC also suppresses the mIPSC frequency. **a.** Experimental design for mitral cell-specific dystroglycan deletions in the OB using retro-AAVs encoding a control gRNA or the dystroglycan (*Dag1*) gRNA together with tdTomato that are injected into the piriform cortex of constitutively CAS9 expressing mice. The retro-AAVs infect axons projecting from OB mitral/tufted cells and thereby delete dystroglycan specifically in mitral/tufted cells of the OB. **b.** Representative fluorescence images of OB sections stained for gephyrin as an inhibitory synapse marker (green) and tdTomato expressed by the retro-AAVs (red). **c** & **d**. Dystroglycan deletion *in vivo* does not change the density or size of gephyrin-positive synaptic puncta in the OB. Sections from mice infected as shown in a and b were analyzed by quantitative immunocytochemistry for the postsynaptic inhibitory synapse marker gephyrin (green) (c, sample images; d, summary graphs of puncta densities (top) and size (bottom)). Only gephyrin puncta that co-localized with tdTomato-expressing mitral cell somata and dendritic segments were quantified. Puncta densities and sizes are plotted as analyzed per animal, not per region-of-interest. **e**. Representative fluorescence image of a mitral cell filled with neurobiotin (blue) via the patch pipette in an OB section from a mouse with CRISPR-induced dystroglycan (*Dag1*) deletion in mitral cells (tdTomato expressed by the retro-AAVs is shown in red). **f**-**h**. The dystroglycan (*Dag1*) deletion decreases the mIPSC frequency *in vivo* in mitral cells (f, representative mIPSC traces recorded in the presence of TTX; g, cumulative probability plots of the interevent interval and summary graph of the mIPSC frequency; h, cumulative probability plots and summary graph of the mIPSC amplitudes). **i**-**k**. The dystroglycan (*Dag1*) deletion greatly decreases the amplitude of IPSCs at GC→MC synapses evoked by extracellular stimulation, as documented by input/output curves to control for possible variations in stimulating electrode placement (i, representative IPSC traces; j, summary plot of input/output amplitude measurements; k, summary graph of the slope of the input/output curves). **l**. The mitral cell-specific dystroglycan (*Dag1*) deletion causes a massive increase in the coefficient of variation of evoked IPSCs at GC→MC synapses, suggesting a decrease in release probability. **m** & **n**. Consistent with a decreased release probability, the dystroglycan (*Dag1*) deletion induces a large increase in the paired-pulse ratio (m, representative traces; n, summary plot of the paired-pulse ratio). **o**-**r**. Dystroglycan deletion in the mPFC in vivo decreases the mIPSC frequency monitored in Layer 5 pyramidal neurons (o, representative image of an mPFC section; p, representative mIPSC traces recorded in the presence of TTX; q, cumulative probability plots of the interevent interval and summary graph of the mIPSC frequency; r, cumulative probability plots and summary graph of the mIPSC amplitudes). Numerical data are means ± SEM; n’s (cells/experiments) are indicated in the summary graph bars (d) or above the sample traces (f, i, m, and p, and apply to all graphs in an experimental series. Statistical analyses were performed using Student’s t-test in d, g, h, k, l, q, r, and by two-way ANOVA in j & n, with * = p<0.05, ** = p<0.01, *** = p<0.001, and *** = p<0.0001.

Dystroglycan is expressed not only by neurons, but also by astrocytes and pericytes, both of which target dystroglycan to the basal lamina encapsulating blood vessels in brain (Fig. S7). We next determined whether dystroglycan acts as a postsynaptic ligand in mitral cells to Neurexin-3 expressed by granule cells. To achieve a specifically postsynaptic deletion of dystroglycan in mitral cells, we crossed CAS9 conditional knockin mice^55^ with tBet-Cre mice, resulting in expression of CAS9 only in mitral/tufted cells. Infection of the OB with AAVs expressing the dystroglycan gRNA and Cre-dependent DIO-tdTomato then causes a selective dystroglycan deletion in mitral/tufted cells, with infected mitral cells visualized via their tdTomato expression (Fig. 7e). This enabled us to performed whole-cell patch-clamp recordings from infected mitral cells in which dystroglycan had been deleted.

We found that the dystroglycan deletion produced a large decrease (∼40%) in mIPSC frequency, and a smaller decrease in mIPSC amplitude, without a change in intrinsic electrical properties or mIPSC kinetics (Fig. 7f-7h, S7d-S7f). Moreover, the dystroglycan deletion induced a comparable decrease (∼40%) in the amplitude of evoked GC→MC IPSCs (Fig. 7i-7k), again without a change in kinetics (Fig. S7g). This decrease in IPSC amplitude was accompanied by a large increase (∼80%) in the coefficient of variation of evoked IPSCs, and by an inversion of paired-pulse suppression to paired-pulse facilitation at short interstimulus intervals (Fig. 7m, 7n).

The results of the dystroglycan deletion experiments are unexpected in that it was previously argued that dystroglycan is important for the formation and not the operation of a subset of GABAergic synapses, and that its synaptic function does not involve binding to neurexins^34, 56^. Therefore we aimed to validate these results in a second set of experiments in which we specifically deleted dystroglycan in mitral cells by infecting the piriform cortex of CAS9 conditional knockin mice with retro-AAVs expressing dystroglycan or control gRNAs and tdTomato (Fig. 8a, 8b). The retro-AAVs are taken up by axonal projections from the mitral cells to the piriform cortex, resulting in the selective deletion of dystroglycan from mitral cells in the OB without any stereotactic injections of the OB.

Again, the dystroglycan deletion in mitral cells had no apparent effect on inhibitory synapse density or size on mitral cells as examined using immunocytochemistry for the inhibitory synapse marker gephyrin (Fig. 8c, 8d). The postsynaptic mitral cell deletion of dystroglycan, however, did cause a pronounced functional impairment. Patch-clamp recordings from mitral cells uncovered a robust decrease (∼40%) in mIPSC frequency but not amplitude (Fig. 8e-8h). No change in intrinsic electrical properties or mIPSC kinetics were present (Fig. S8a-S8c), and OB slices from male and female mice exhibited similar decreases in mIPSC frequency (Fig. S8d). Moreover, the dystroglycan deletion greatly decreased (∼50%) the amplitude of evoked GC→MC IPSCs (Fig. 8i-8k), and increased (∼100%) the coefficient of variation of IPSCs without changing the kinetics of the iPSCs (Fig. 8l, S8e). Consistent with this result suggesting a decrease in release probability, the dystroglycan deletion also converted paired-pulse responses from depressed to facilitated at short interstimulus intervals (Fig. 8m, 8n).

Viewed together, the dystroglycan deletion phenotype is a mirror image of the *Nrxn3* KO phenotype, with a dramatic loss of GC→MC synaptic strength due to a decrease in release probability but without detectable decrease in synapse numbers. These results, consistent with the rescue of the Nrxn3 KO phenotype only with Nrxn3α splice variants that bind to dystroglycan strongly support the notion that Nrxn3α enables GC→MC synaptic function via binding to dystroglycan. As a final question we thus asked whether such a mechanism also applies to mPFC role of *Nrxn3*. Indeed, when we applied the CRISPR-mediated deletion of dystroglycan to the mPFC, we also detected a significant decrease in mIPSC frequency without a change in mIPSC amplitude (Fig. 8o-8r). Moreover, no major changes in the intrinsic electrical properties or mIPSC kinetics were observed (Fig. S8f-S8h). Overall, these data support the notion that *Nrxn3* also shapes a subset of inhibitory synapses in the mPFC by binding to dystroglycan.

## SUMMARY

Here we show that binding of presynaptic Nrxn3α to postsynaptic dystroglycan organizes the functional architecture of inhibitory GC→MC synapses in the OB and of inhibitory synapse on layer 5/6 neurons in the mPFC. We demonstrate that the Nrxn3α/ dystroglycan interaction is not essential for the formation of these synapses, but renders these synapses competent for neurotransmitter release by enabling a normal release probability. Moreover, we find that the role of Nrxn3α at these synapses is controlled by a combinatorial code of alternative splicing whereby SS2 and SS4 of Nrxn3α collaborate to determine the release probability as an ‘AND/OR’ logic gate. Thus, our data propose a feedback mechanism by which binding of presynaptic Nrxn3α to postsynaptic dystroglycan is required in order to enable Nrxn3α to organize the presynaptic neurotransmitter release machinery. The evidence for these overall conclusions can be summarized as follows:

First, deletion of Nrxn3α lowered the strength of inhibitory GC→MC synapses by more than a half; this impairment was rescued by Nrxn3α but not by Nrxn3β, with Nrxn3α only being active when its alternatively spliced SS4 and/or SS2 sites contain no insert (Fig. 1, 2; S1-S4). SS2 is dominant in this combinatorial splice code because even when SS4 is spliced out, the longer insert in SS2 (SS2ab) blocked the function of Nrxn3α (Fig. 1). In granule cells in vivo, nearly 100% of *Nrxn3* mRNAs contain an insert in SS4 and more than 90% of mRNAs lack in insert in SS2, suggesting that a Nrxn3α/dystroglycan complex is normally formed (Fig. S3, S4). However, it is unknown whether Nrxn3-SS2 and -SS4 alternative splicing is activity-dependent in these neurons, and this ratio might change during specific behavioral states or during maturation of adult-born OB granule cells, which could regulate GC→MC synaptic transmission by altering the Nrxn3α/dystroglycan interaction.

Second, the mechanism by which deletion of Nrxn3α suppressed GC→MC synaptic transmission consisted of a decrease in the presynaptic release probability, as shown by an increased coefficient of variation of IPSCs, a dramatic shift in paired-pulse ratio, and a lack of change in synapse numbers (Fig. 3-4). Thus, this phenotype is similar to the phenotype previously observed in neurexin-deficient other synapses in which a disorganization of calcium channels impairs the coupling of voltage-gated calcium influx to neurotransmitter release^5–7^.

Third, the *Nrxn3* KO phenotype at GC→MC synapses is fully rescued by a construct that contains only the LNS2-domain of the extracellular LNS- and EGF-domains of Nrxn3α, provided the LNS2-domain lacks an insert in SS2 (Fig. 2, 4). This rescue was observed both in cultured neurons and in vivo, suggesting that even though the *Nrxn3* deletion causes a decrease in release probability of its resident nerve terminal, a trans-synaptic interaction of Nrxn3α with a postsynaptic trans-ligand is required for GC→MC synapse function. Notably, these findings support a “Swiss Army Knife”-like functional modularity of α-Neurexins, with their large size and presence of an array of independent binding units endowing them with the ability to simultaneously engage diverse trans-synaptic ligands in orchestrating synapse properties.

Fourth, the CRISPRi-mediated inhibition of expression and CRISPR-mediated deletion of dystroglycan in postsynaptic mitral cells caused the same phenotype as the presynaptic Nrxn3 deletion in cultured neurons and in vivo (Fig. 6-8). Since this is the first validation of the physiological relevance of neurexin-binding to dystroglycan, which was previously questioned^34^, we aimed to confirm this conclusion using two different CRISPR-approaches to delete dystroglycan in vivo from mitral cells, namely direct infection of the OB with AAVs expressing the dystroglycan-specific guide-RNA only in mitral cells (Fig. 7), and retrograde infection of only mitral cells by administration of retro-AAVs expressing the guide-RNA into the piriform cortex (Fig. 8). Importantly, the postsynaptic dystroglycan deletion had no effect on synapse numbers in vivo, but caused the same increase in the coefficient of variation of IPSCs and in their paired-pulse ratio as the presynaptic Nrxn3 deletion. Thus, our findings indicate that postsynaptic dystroglycan binding to presynaptic Nrxn3α retrogradely regulates the presynaptic release probability without affecting synapse formation as such. The strongest evidence for this conclusion comes from the selective rescue of the *Nrxn3* deletion phenotype by Nrxn3α constructs still capable of interacting with dystroglycan as shown previously^26^.

Fifth, deletion of *Nrxn3* or of dystroglycan from mPFC neurons produced the same overall phenotype as these deletions induced in OB neurons, namely a loss of inhibitory synaptic strength associated with a change in release probability (Fig. 6, 8). Most importantly, the *Nrxn3* deletion phenotype was again completely rescued by the LNS2-only Nrxn3α construct lacking an insert in SS2 (Fig. 6). The observed phenotype in the mPFC was not as severe as that found in the OB. This presumably derives from the fact that we analyzed a relatively homogeneous population of inhibitory GC→MC synapses in the OB in which Nrxn3α/dystroglycan binding invariably shapes synapse properties, where we examined a heterogeneous mixture of distinct inhibitory synapses in the mPFC in which only some synapses may utilize the Nrxn3α/dystroglycan signaling mechanism.

Previous work demonstrating that dystroglycan is important for synapses is generally consistent with our results, but most of these studies found a role in synapse formation instead of synapse function^34, 40, 56^. Since the previous studies were performed in the hippocampus and somatosensory cortex, while we examined the OB and mPFC, it is possible that the results are due to differences in the type of synapses studied. Früh et al. (2016) also concluded that the function of dystroglycan in synapses is independent of neurexins, which is plausible since it is a different brain region compared to that studied here, although neurexins and their binding to the dystroglycan mutant used in the hippocampal studies were not actually examined. Alternatively, dystroglycan may separately regulate the initial targeting of CCK+ interneurons, which may be the proximal cause of fewer CCK+ synapses in dystroglycan KO mice where dystroglycan was depleted during development^34, 40^. At these synapses and others, only once synapses have formed might signaling between dystroglycan and Nrxn3α become critical for sustaining presynaptic release.

Arguably, our most surprising result is that trans-synaptic binding of presynaptic Nrxn3α to postsynaptic dystroglycan is required for the ability of Nrxn3α to organize a fully functional presynaptic release machinery. What is the nature of the dystroglycan-activated signal in presynaptic terminals – is it a rearrangement of Nrxn3α (e.g., dimerization), or an independent additional signal? This question is likely not only important for understanding the functional molecular architecture of synapses, but also for insight into how mutations in genes associated with dystroglycan, such as mutations in the glycosylating enzymes for dystroglycan or their cytoplasmic binding proteins, and in Nrxn3α produce neurodevelopmental disorders^41–43^. Our findings define the core interaction of Nrxn3α with dystroglycan as functionally essential for inhibitory synapses in at least two brain regions, but they do not yet reveal the detailed molecular signaling that organizes the presynaptic release machinery, a question that will need to be addressed in future.

In summary, we have defined a trans-synaptic signaling complex that performs an indispensable role in organizing the presynaptic release probability at a subset of inhibitory synapses, which compared to excitatory synapses are poorly understood. Our findings underscore the notion that individual functional properties of diverse synapses must be systematically studied at a molecular level because information processing in the brain not only depends on how neural circuits are wired via synaptic connections, but also on the functional properties of these connections. Moreover, our current findings add to our understanding of the diverse synaptic roles of neurexins. One might ask, however, why the organization of synapses is so complicated, and why neurexins perform so many diverse functions in different types of synapses. This more philosophical question is part of the larger issue of why the brain needs to have so many different types of neurons and synapses to operate properly. Naturally, this question is unanswerable at present, but it is striking that with neurexins, a single gene family is used to diversify different types of synapses in the context of distinct circuits. Instead of expressing possibly hundreds of genes to determine synapse identity, with the neurexins the brain expresses only three genes, whose products are uniquely capable of generating thousands of isoforms and interacting with dozens of ligands, to endow different synapses with distinct properties – a remarkable simplification of the mechanism of synapse diversification. In this view, neurexins do not complicate the design of synapses, but simplify it, even though the overall need for diversity creates a panoply of different molecular pathways whose full extent remains to be characterized.

## MATERIALS AND METHODS

### Animals

Nrxn3 conditional knockout (cKO) mice were generated as described. Other mouse lines used in this paper include: tBet-Cre^57^, constitutive cas9-knockin (KI) (Jax, 024858), conditional cas9-KI (Jax, 024857), vGAT-Cre (Jax, 028862) and RiboTag mice (Jax, stock# 029977). For analyzing mitral/tufted cell–specific or inhibitory neuron (primarily granule cell) translating mRNA, RiboTag mice were crossed with hemizygous tBet-Cre mice^57^ and hemizygous vGAT-Cre mice, respectively. For all experiments using constitutive and conditional Cas9-KI mice, mice were maintained at homozygosity for the KI alleles. For mitral/tufted cell–specific deletion of *Dag1*, conditional cas9-KI mice were crossed with mice carrying the tBet-Cre allele. Only mice with a single allele of Cre were used for experiments. All mice were weaned at 20 days of age and housed in groups of 2 to 5 on a 12 hr light/dark cycle with access to food and water *ad libidum*. All procedures conformed to National Institutes of Health Guidelines for the Care and Use of Laboratory Mice and were approved by the Stanford Animal Use Committees [Administrative Panel for Laboratory Animal Care (APLAC/) Institutional Animal Care and Use Committee (IACUC)].

### Plasmids

Lentiviral vectors for expression of Cre and ΔCre (truncated, inactive) recombinase driven by the human synapsin-1 promoter (i.e. have been described previously^58^. For all other experiments using the lentiviral backbone with a human synapsin-1 vector (i.e. FSW), an empty vector was used as a control. For all Nrxn3 rescue constructs, a single HA tag was positioned between the native signal peptide and was flanked by linker sequences (i.e. glycine-glycine-serine upstream and glycine-serine downstream). All culture rescue constructs were incorporated into the FSW lentiviral backbone. A library of previously published cDNA’s^12, 44^ were used to clone Nrxn3alpha and Nrxn3beta splice variants described in Figs. 1-2. For all truncation constructs (Fig. 2), that lacked LNS6, upstream domains were fused at the same position that LNS6 would normally be, thus preserving the downstream stalk region, transmembrane domain, and cytoplasmic sequence.

The adeno-associated virus (AAV) serotypes used in this study were AAV-DJ and rAAV2-retro for retrograde experiments^59^. AAV backbones were generated to allow the expression of expression of Cre and ΔCre fused to tdTomato, minimal Nrxn3 LNS2 rescues (with and without SS2), and gRNA’s targeting *Dag1* with soluble tdTomato driven by the hSynI promoter in a Cre-sensitive (i.e. with DIO) or constitutive manner. For CRISPRi lentiviral backbones, a scrambled gRNA control was generated (5’-GCGCCAAACGTGCCCTGACG-3’). For targeting of dystroglycan several gRNA’s were initially screened. The final gRNA that performed best following a functional screen was 5’-AGCTTCGCGCGGAGTCCCCG-3’. CRISPRi was performed using a lentiviral backbone described previously^60^, with expression of gRNA driven by the U6 promoter and the inactive Cas9 fused to KRAB driven by the EFS promoter.

For *in vivo* CRISPR experiments, two gRNA were used to ensure efficient targeting of *Dag1* including one driven by a U6 promoter (i.e. 5’-tggttaggttctcccccacg-3’) and another by a H1 promoter (i.e. 5’-accgtggttggcattccaga-3’). These gRNA were published previously^39^. Scrambled gRNA sequences were used as controls.

### Primary Antibodies

The following antibodies were used at the indicated concentrations (IHC-immunohistochemistry; ICC-immunocytochemistry): anti-Dystroglycan rabbit (Abcam Cat# ab199768; 1:250 IHC), anti-HA rabbit (Cell Signaling Cat# 3724; 1:500 IHC), anti-Gephyrin mouse (Synaptic Systems Cat# 147 011; 1:1000 ICC), anti-Gephyrin guinea pig (Synaptic Systems Cat# 147 318; 1:250 IHC), anti-Homer1 rabbit (Synaptic Systems Cat# 160 003, 1:1000), anti-MAP2 chicken (Encorbio Cat# CPCA-MAP2; 1:1000), anti-GABA_A_Rα1 (Synaptic Systems Cat# 224 203; 1:250 live ICC), anti-GABA_A_Rα2 (Synaptic Systems Cat# 224 103; 1:250 live ICC), anti-GABA_A_Rγ2 (Synaptic Systems Cat# 224 003; 1:250 live ICC), anti-Synaptophysin-2 rabbit (homemade, Wang et al., 2021; 1:500), and anti-vGAT guinea pig (Synaptic Systems Cat# 131 004; 1:1000 ICC).

### Cell Culture

#### Primary neuron cultures (containing glia)

Hippocampal, olfactory bulb, and cortical neurons were cultured from newborn mice as described previously^45, 61^ with some modifications. Tissue was dissected and mixed regardless of gender. In general, pooling tissue from three to six mice in a given preparation was used to generate cultures. Dissected hippocampi, olfactory bulbs, or cortices were digested for 20 min with 10 U/ml papain in Hank’s buffered saline (HBS) in an incubator, washed with HBS, dissociated in plating media (MEM supplemented with 0.5% glucose, 0.02% NaHCO_3_, 0.1 mg/ml transferrin, 10% FBS, 2 mM L-glutamine, and 0.025 mg/ml insulin), and seeded on Matrigel (BD Biosciences) precoated coverslips placed inside 24-well dishes. For olfactory bulb neurons, the day after plating, 95% of media was replaced with MEM (GIBCO) supplemented with 2% B27 (GIBCO), 0.5% w/v glucose, 100 mg/l transferrin, 5% fetal bovine serum. For hippocampal and cortical neurons, the day after plating, 95% of the plating medium was replaced with neuronal growth medium lacking serum (Neurobasal-A medium supplemented with 2% B27 supplement and 0.5 mM L-glutamine). At DIV2–3 (for hippocampal and olfactory bulb cultures) or DIV3–4 (for cortical cultures), 50% of the medium was exchanged with fresh growth medium additionally supplemented with 4 µM AraC (Sigma-Aldrich) to restrict glial overgrowth. When applicable, neurons were infected between DIV3-4 with lentiviruses expressing EGFP-tagged ΔCre (control) or Cre without and/or with the indicated rescue constructs driven by the synapsin promoter. For long-term culture of hippocampal and cortical neurons, 25% fresh media was added every 4–5 d starting from DIV7. A partial media change was performed only once on DIV7 for OB neurons to preserve cell health.

#### HEK293T cells

HEK293T cells (ATCC CRL-11268) were grown in complete DMEM (cDMEM), which consisted of DMEM (Gibco), 5% FBS (Sigma), penicillin, and streptomycin. All transfections were performed using lipofectamine 3000 (Invitrogen). For co-culture assays, HEK293T cells were plated on 12-well plates and transfected at ∼90% confluency according to manufacturer’s instructions.

### Preparation of Viral Particles

#### AAV preparation

The adeno-associated virus (AAV) serotypes used in this study were AAV-DJ and rAAV2-retro for retrograde experiments^59^. HEK293T cells were transfected with the helper plasmid, the serotype-specific plasmid and the AAV plasmid using homemade calcium phosphate solution. Cells were dissociated and precipitated 72 hours post-transfection. Nuclei were lysed by three times of freeze-thaw cycles and were later treated with Benzonase nuclease (Sigma-Aldrich, cat # E1014). The supernatant then underwent iodixanol gradient ultracentrifugation for 3 hours at 65,000 rpm at 4LC in a S80AT3 rotor. AAV were then concentrated using filtered centrifugation and dialyzed in minimal essential media (MEM).

#### Lentivirus preparation

Recombinant lentiviral particles were produced in HEK293T cells by co-transfecting cells with long terminal repeat (LTR) containing vector and helper plasmids (pRSV-REV, pMDLg/gRRE, and pVSVG) using calcium phosphate. Media were exchanged 1 h before transfection and included 25 µM chloroquine diphosphate. Per 75 cm2 of cells, 0.5 ml of 250 mM CaCl2 containing molar equivalents of DNA (12 µg of LTR-containing vector, 3.9 µg pRSV-REV, 8.1 µg pMDLg/gRRE, and 6.0 µg pVSVG) was added dropwise to an equal volume of 2X-HBS (0.4 M NaCl, 10 mM KCl, 1.5 mM Na2HPO4, 0.2% glucose, and 38.4 mM HEPES, pH 7.05) under vigorous mixing, incubated for 20 min at room temperature, and added dropwise to the cells. 16–20 h following transfection, cells were washed with plain DMEM and replaced with neuronal growth media lacking AraC. After 24 h, media containing lentiviral particles were cleared by centrifugation (1,500 ×g, 10 min), aliquoted, and snap-frozen. Neuronal cultures were infected with lentivirus on DIV3 or 4 by adding 25–30 µl of viral supernatant per well of a 24-well plate.

### Electrophysiology

#### Culture electrophysiology

Cultured neurons were collected and recorded at DIV 14-17. Electrophysiology recordings were performed at room temperature, performed in whole-cell patch-clamp mode using concentric extracellular stimulation electrodes. The glass pipettes (2-3 MΩ filled with intracellular pipette solution) were pulled from borosilicate glass capillaries with a vertical micropipette puller (PC-10, Narishige). After formation of the whole-cell configuration and equilibration of the intracellular pipette solution, the series resistance was adjusted to 8–10 MΩ. Synaptic currents were monitored with a Multiclamp 700B amplifier (Molecular Devices). A bipolar stimulation electrode (FHC, Bowdoinham, ME) was placed 100-150 µm from the soma of the neurons recorded to apply focal square pulse stimuli (duration 1 ms) and trigger evoked synaptic responses. The frequency, duration, and magnitude of the extracellular stimulus were controlled with a Model 2100 Isolated Pulse Stimulator (A-M Systems) synchronized with Clampex 9 data acquisition software (Molecular Devices). The whole-cell pipette solution contained (in mM): 120 CsCl, 5 NaCl, 1 MgCl_2_, 10 HEPES, 10 EGTA, 0.3 Na-GTP, 3 Mg-ATP, and 5 QX-314 (pH 7.2, adjusted with CsOH). The bath solution contained (in mM): 140 NaCl, 5 KCl, 2 MgCl_2_, 2 CaCl_2_, 10 HEPES, and 10 glucose (pH 7.4, adjusted with NaOH). IPSCs, as well as AMPAR- or NMDAR-mediated EPSCs, were pharmacologically isolated by adding blockers against AMPA receptor (CNQX, 10 μM), NMDA receptor (APV, 50 μM), or GABA_A_ receptor (picrotoxin, 50 μM) to the extracellular solution. Spontaneous mIPSCs and mEPSCs were monitored in the presence of tetrodotoxin (TTX, 1 μM) to block action potentials. Miniature events were analyzed in Clampfit 9 (Molecular Devices) using the template matching search and a minimal threshold of 5 pA and each event was visually inspected for inclusion or rejection by an experimenter blind to the recording condition.

#### Slice Electrophysiology

Two to three weeks after viral injection, mice were anesthetized via isoflurane inhalation and brains were quickly dissected. The dissected brain was sliced in ice-cold oxygenated (95% O_2_ and 5% CO_2_) cutting solution (228 mM sucrose, 11 mM glucose, 26 mM NaHCO_3_, 1 mM NaH_2_PO_4_, 2.5 mM KCl, 7 mM MgSO_4_, and 0.5 mM CaCl_2_). Horizontal sections for OB and coronal sections for mPFC, both of which were 300 µm thick, were obtained by using a vibratome. Slices were quickly transferred to oxygenated artificial cerebrospinal fluid (ACSF; 119 mM NaCl, 2.5 mM KCl, 1 mM NaH_2_PO_4_, 1.3 mM MgSO_4_, 26 mM NaHCO_3_, 10 mM glucose, and 2.5 mM CaCl_2_) at 32°C for 30 min. Slices were allowed to recover at room temperature for an additional 30 min. The recording chamber was temperature controlled and set to 32°C, and ACSF was perfused at 1 mL/min. The internal solution for whole-cell patch clamp contained 135 mM CsCl, 10 mM HEPES, 1 mM EGTA, 1 mM Na-GTP and 4 mM Mg-ATP pHed to 7.4. 10mM QX314-bromide was added for evoked recordings. 0.2% neurobiotin (VectorLab) was included for morphological reconstruction. The pipette resistance ranged from 1.8 to 2.5 MΩ. Mitral cells were identified in the mitral cell layer in the OB and mPFC neurons were identified either by fluorescent reporter or pyramidal-shaped neuron in deep layer. Access resistance was under 10 MΩ (for mitral cells) and 15 MΩ for mPFC neurons throughout the experiment. 1µM TTX (Tocris), 20µM CNQX (Tocris) and 50µM D-AP5 (Tocris) were included in the bath for mIPSC. 20µM CNQX (Tocris) and 50µM D-AP5 (Tocris) were included in the bath for evoked IPSC recordings. All recordings were done in voltage-clamp mode with the holding potential of −70mV. For eIPSC stimulation, concentric bipolar electrode was used. For GC→MC eIPSC, the stimulating electrode was placed directly below the mitral cell with constant distance roughly at the junction between internal plexiform layer and granule cell layer, 30 µm below the surface of the slice. For eIPSC in mPFC, the stimulating electrode was placed parallel to the recorded cell in the same layer with constant distance and 30 µm below the surface of the slice. The experimenter was blind to the treatment groups during recordings and analysis.

#### Stereotactic injections

Mice were prepared for stereotactic injections using standard procedures approved by the Stanford University Administrative Panel on Laboratory Animal Care. Mice were anesthetized by 0.2 mL avertin working solution per 10 grams body weight. The avertin stock solution was made by dissolving 5 grams tribromoethanol into 5 mL T-amyl alcohol, which was further diluted 80 folds in DPBS to make the avertin working solution. The coordinates (AP/ML/DV from Bregma) and volumes for the intercranial injections are (1) +4.3/±0.85/-1.7 and +5.3/±0.6/-1.5 with 1.0 uL virus for the olfactory bulbs and (2) −0.7/±3.7/-4.75 with 0.75 uL virus for the piriform cortex (for retrograde targeting of mitral cells). For injection into the piriform cortex, the mouse brains were aligned to have less than 0.05mm difference on the DV axis at −1.00 (AP) between ±3.00 (ML) positions. The reference zero point for DV is on the surface of the OB for OB injection and the surface of the skull at AP/ML of −0.7/0.0 for piriform cortex injection.

### Purification of Tissue mRNA

Wild-type (CD1) mice at 8 weeks of age were euthanized using isoflurane and decapitated. Several brains regions including the cortex, olfactory bulb, cerebellum, hippocampus, and pons/medulla were quickly dissected and snap-frozen in liquid nitrogen or dry ice and transferred to −80°C storage until processing. The specimen was subjected to RNA extraction using the QIAGEN RNeasy Micro kit.

### Purification of Ribosome-bound mRNA

RiboTag mice^62^ were crossed with tBet-Cre mice^57^ or vGAT-Cre mice. After OB extraction described above, the frozen bulbs were partially thawed in fresh homogenization buffer at 10% (w/v) and Dounce homogenized. Homogenates underwent centrifugation, and 10% of the supernatant was used as input. The remaining supernatant was incubated with prewashed anti-HA magnetic beads (Thermo Fisher Scientific) overnight at 4°C. The beads were washed three times with a high-salt buffer followed by elution with RLT lysis buffer containing 2-mercaptoethanol. The sample and the input were then subjected to mRNA extraction described above. RNA concentration was determined using a NanoDrop 1000 Spectrophotometer (Thermo) and stored at −80°C until downstream analysis.

### qPCR

Quantitative reverse transcription (RT)–PCR was performed in triplicates for each condition with QuantStudio 3 (Thermo Fisher Scientific). RNA (20 ng) was used each reaction, in conjunction with TaqMan Fast Virus 1-Step Master Mix (Thermo Fisher Scientific) and gene-specific qRT-PCR probes [IDT (integrated DNA technologies)] as described previously (66). Predesigned PrimeTime qPCR probe assays (IDT) were used for vGluT1 (Mm.PT.58.12116555), vGaT (Mm.PT.58.6658400), aquaporin-4 (Mm.PT.58.9080805), MBP (Mm.PT.58.28532164), ActB (Mm.PT.39a.22214843.g), Cbln1 (Mm.PT.58.12172339), Cbln2 (Mm.PT.58.5608729), Cbln4 (Mm.PT.58.17207498), Grid1 (Mm.PT.58.32947175), and Grid2 (Mm.PT.58.12083939), Nlgn1 (Mm.PT.58.30240881), Nlgn2 (Mm.PT.58.16799702), Nlgn3 (Mm.PT.58.31138258, Nxph1 (Mm.PT.58.13767897), Nxph2 (Mm.PT.58.28481365), Nxph3 (Mm.PT.12688150), Nxph4 (Mm.PT.58.11246838), LRRTM1 (Mm.PT.58.42587284.g) LRRTM2 (Mm.PT.58.6337058.g), LRRTM3 (Mm.PT.58.31131475), LRRTM4 (Mm.PT.58.11146838), Fam19a1 (Mm.PT.56a.6079538), Fam19a2 (Mm.PT.58.7298614), Fam19a4 (Mm.PT.56a.9330679), Car10 (Mm.PT.58.11765793), Car11 (Mm.PT.58.32895602), Dag1-ex1/ex2 (Mm.PT.58.46076316), Dag1-ex3/ex4 (Mm.PT.58.45967735), and Dag1-ex4/ex5 (Mm.PT.58.5524327). Customed PrimTime qPCR probe assays (IDT) were used for Nrxn1α (forward: TTCAAGTCCACAGATGCCAG; reverse: CAACACAAATCACTGCGGG; probe: TGCCAAAACTGGTCCATGCCAAAG), Nrxn1β (forward: CCTGTCTGCTCGTGTACTG; reverse: TTGCAATCTACAGGTCACCAG; probe: AGATATATGTTGTCCCAGCGTGTCCG), Nrxn1γ (forward: GCCAGACAGACATGGATATGAG; reverse: GTCAATGTCCTCATCGTCACT; probe: ACAGATGACATCCTTGTGGCCTCG), Nrxn2α (forward: GTCAGCAACAACTTCATGGG; reverse: AGCCACATCCTCACAACG; probe: CTTCATCTTCGGGTCCCCTTCCT), Nrxn2β (forward: CCACCACTTCCACAGCAAG; reverse: CTGGTGTGTGCTGAAGCCTA; probe: GGACCACATACAT CTTCGGG), Nrxn3α (forward: GGGAGAACCTGCGAAAGAG; reverse: ATGAAGCGGAAGGACACATC; probe: CTGCCGTCATAGCTCAGGATAGATGC), Nrxn3β (forward: CACCACTCTGTGCCTATTTC; reverse: GGCCAGGTATAGAGGATGA; probe: TCTATCGCTCCCCTGTTTCC), Nlgn1 (forward: GGTTGGGTTTGGTATGGATGA; reverse: GATGTTGAGTGCAGTAGTAATGAC; probe: TGAGGAACTGGTTGATTTGGGTCACC), Nlgn2 (forward: CCGTGTAGAAACAGCATGACC; reverse: TGCCTGTACCTCAACCTCTA; probe: TCAATCCGCCAGACACAGATATCCG), and Nlgn3 (forward: CACTGTCTCGGATGTCTTCA; reverse: CCTCTATCTGAATGTGTATGTGC; probe: CCTGTTTCTTAGCGCCGGATCCAT).

Assays generating C_t_ values >35 were omitted. C_t_ values for technical replicates (duplicate or triplicate) differed by less than 0.5. C_t_ values were averaged for technical replicates. Data were normalized to the arithmetic mean of *ActB* and *Gapdh* using the 2^-ΔΔCt^ method.

### Junction-flanking PCR

The following primers anneal to constitutive exon sequences that flank splice junctions and thus amplify *Nrxn1-3* mRNA transcripts with or without alternative splice sequences (splice site, forward primer, reverse primer): Nrxn1 SS2 (5’-TGGGATCAGGGGCCTTTGAAGCA-3’, 5’-GAAGGTCGGCTGTGCTGGGG-3’), Nrxn1 SS4 (5’-CTGGCCAGTTATCGAACGCT-3’, 5’-GCGATGTTGGCATCGTTCTC-3’), Nrxn2 SS2 (5’-GCACGACGTCCGGGTTACCC-3’, 5’-GGTCGGCTGTGTTGGGGCTG-3’), Nrxn2 SS4 (5’-CAACGAGAGGTACCCGGC-3’, 5’-TACTAGCCGTAGGTGGCCTT-3’), Nrxn3 SS2 (5’-TCCGGGGCCTTTGAGGCCAT-3’, 5’-GCGGTACTTGGGCTTCCACCA-3’), Nrxn3 SS4 (5’-CCAGGAATGGGGGAAATGCT-3’, 5’-TTGTCCTTTCCTCCGATGGC-3’). cDNA was synthesized from equal amounts of 1) adult brain regions, 2) primary neuron/glia culture mRNA, or 3) immunoprecipitated mRNA from mitral/tufted cells or granule cells and total input mRNA from the OB. Junction-flanking PCR was then performed with equal amount of cDNA from groups being compared. The PCR products were separated on homemade MetaPhor agarose gel (Lonza) and stained with GelRed. Stained gel was imaged at subsaturation using the ChemiDoc Gel Imaging System (Bio-Rad). Quantification was performed using Image Lab (Bio-Rad) or ImageStudioLite (LI-COR). Intensity values were normalized to the size of DNA products to control for intensity differences caused by different dye incorporation owing to varied DNA length.

### Immunocytochemistry

For live surface-labeling experiments, primary neurons were first washed at room temperature once with a HEPES bath solution, which contained the following (in mM): 140–150 NaCl, 4–5 KCl, 2 CaCl_2_, 1 MgCl_2_, 10 glucose, and 10 HEPES, with pH adjusted to 7.4 with NaOH, and osmolarity of 300 mOsm. Cultures were then incubated at room temperature for 20 min with antibodies recognizing the GABA_A_Rα1, GABA_A_Rα2, or GABA_A_Rγ2 diluted in HEPES bath solution. Cultures were then gently washed three times with HEPES bath solution, followed by fixation for 20 min at room temperature with 4% (wt/vol) PFA. Following fixation, cultures were washed three times with Dulbecco’s PBS (DPBS). Cultures were blocked for 1 h at room temperature with antibody dilution buffer (ADB) without Triton X-100(−), which contains 5% normal goat serum diluted in DPBS. Cells were then labeled with Alexa Fluor–conjugated secondary antibodies (1:1,000; Invitrogen) diluted in ADB(−) for 2 h at room temperature. Cultures were then incubated for 10 minutes with 4% PFA for post-fixation and stains proceeded as described above.

For all immunocytochemistry experiments, cells were washed and then permeabilized and blocked for 1 h with ADB with tx-100(+) which contains 0.3% tx-100 and 5% normal goat serum diluted in DPBS. Non-surface primary antibodies were diluted in ADB(+) and cells were incubated in the cold-room overnight or for 2 hrs at RT. Cultures were washed three times with DPBS and then incubated with Alexa Fluor-conjugated secondary antibodies (1:1000; Invitrogen) diluted in ADB(+) for 1 h at RT. After three additional washes, coverslips were inverted onto glass microscope slides with Fluoromount-G mounting media (Southern Biotech).

### Immunohistochemistry

Mice were anesthetized with isoflurane and then transcardially perfused (∼1 ml/min) for 1 min with 0.1M DPBS (RT) followed by 7 min with 4% PFA (Electron Microscopy Services). Olfactory bulbs were dissected and post-fixed for 13 minutes at RT. Tissue was then washed 3 times with DPBS and cryoprotected by a 24-48 hr incubation in 30% sucrose w/v in DPBS. Tissue was embedded in OCT Compound (Sakura), sectioned on the sagittal plane at 30-µm using a cryostat, and stored as floating sections in DPBS. For staining, free-floating sections were incubated with blocking buffer (containing 5% NGS, 1% Pen-Strep, and 0.3% tx-100 in DPBS) for 1 hr at RT. Sections were then incubated with primary antibodies diluted in blocking buffer overnight at RT on a rocker. After 3 washes, sections were incubated with Alexa dye secondary antibodies diluted in blocking buffer for 1-2 hrs at RT. Sections were washed 3-4 times and then mounted on charged glass slides. After drying, sections were dipped in water and allowed to dry again. For confocal imaging, per slide, 4 droplets of Fluormount-G with or without DAPI was added, slides were coverslipped, and nail polish was used to secure the coverslip until mounting medium hardened.

### Confocal Microscopy

All confocal images were acquired at RT using an inverted Nikon A1RSi confocal microscope equipped with a 20x, 60x, or 100x objective (Apo, NA 1.4) and operated by NIS-Elements AR acquisition software. In general, high magnification images were taken at 1,024 × 1,024 pixels with a z-stack distance of 0.3 µm. Low magnification images were taken at 1,024 x 1,024 pixels with Nyquist recommended step size. Line averaging (2X) was used for most images. Images were acquired sequentially in order to avoid bleed-through between channels. Imaging parameters (i.e., laser power, photomultiplier gain, offset, pinhole size, scan speed, etc.) were optimized to prevent pixel saturation and kept constant for all conditions within the same experiment. Images were analyzed using NIS-Elements Advanced Research software (Nikon). For analysis of synapophysin-2 and gephyrin puncta in tissue, local background subtraction was performing using the rolling ball method. For all other image analysis, background was empirically determined and applied equally to all images from aa given imaging session / independent experimental replicate. For quantitative analysis, imaging of brain tissue involved imaging at least 2 regions of interest from 4-5 brain sections. For cultured neurons, two 15-20 µm dendritic segments were analyzed from 8-10 neurons per culture batch, per condition. All ICC/IHC data was collected and analyzed blindly.

### Quantification and Statistical Analysis

Quantifications have been described in the respective materials and methods sections, and statistical details are provided in the figure legends. Statistical significance between various conditions was assessed by determining p-values (95% confidence interval). Statistical analyses were performed using GraphPad Prism 6 software.

For most staining experiments, the “n” represents the average per animal or average per culture. In contrast, for electrophysiology measurements, the “n” represents the total number of cells patched. For biochemical, the “n” generally represents number of animals, independent cultures or pooled samples. Most intergroup comparisons were done by two-tailed Student’s t test or the Welch’s t test. For multiple comparisons, data were analyzed with one- or two-way ANOVA followed by a post-hoc test (e.g. Dunnett’s). Levels of significance were set as * p < 0.05; ** p < 0.01; *** p < 0.001; **** p < 0.0001. All graphs depict means ± SEM.

## ACKNOWLEDGEMENTS

We thank I. Huryeva for excellent technical assistance. This study was supported by grants from the NIMH (MH052804 to T.C.S.; KO1-MH105040-01 to J.H.T), a BBRF Young Investigator Grant (to J.H.T.), and a Stanford Interdisciplinary Graduate Fellowship (SIGF) to (C.Y.W.)

## AUTHOR CONTRIBUTIONS

J.H.T., C.Y.W., and T.C.S. designed, and J.H.T., C.Y.W., and P.Z., conducted all experiments. J.H.T. performed all molecular cloning, RNA analysis, and synapse morphology analysis. C.Y.Z. performed all in vivo recordings and P.Z. performed all in vitro recordings. J.H.T., C.Y.W. and T.C.S. analyzed the data and wrote the manuscript; all authors reviewed and approved the final manuscript.

## EXTENDED DATA FIGURES and LEGENDS

**Figure S1:**
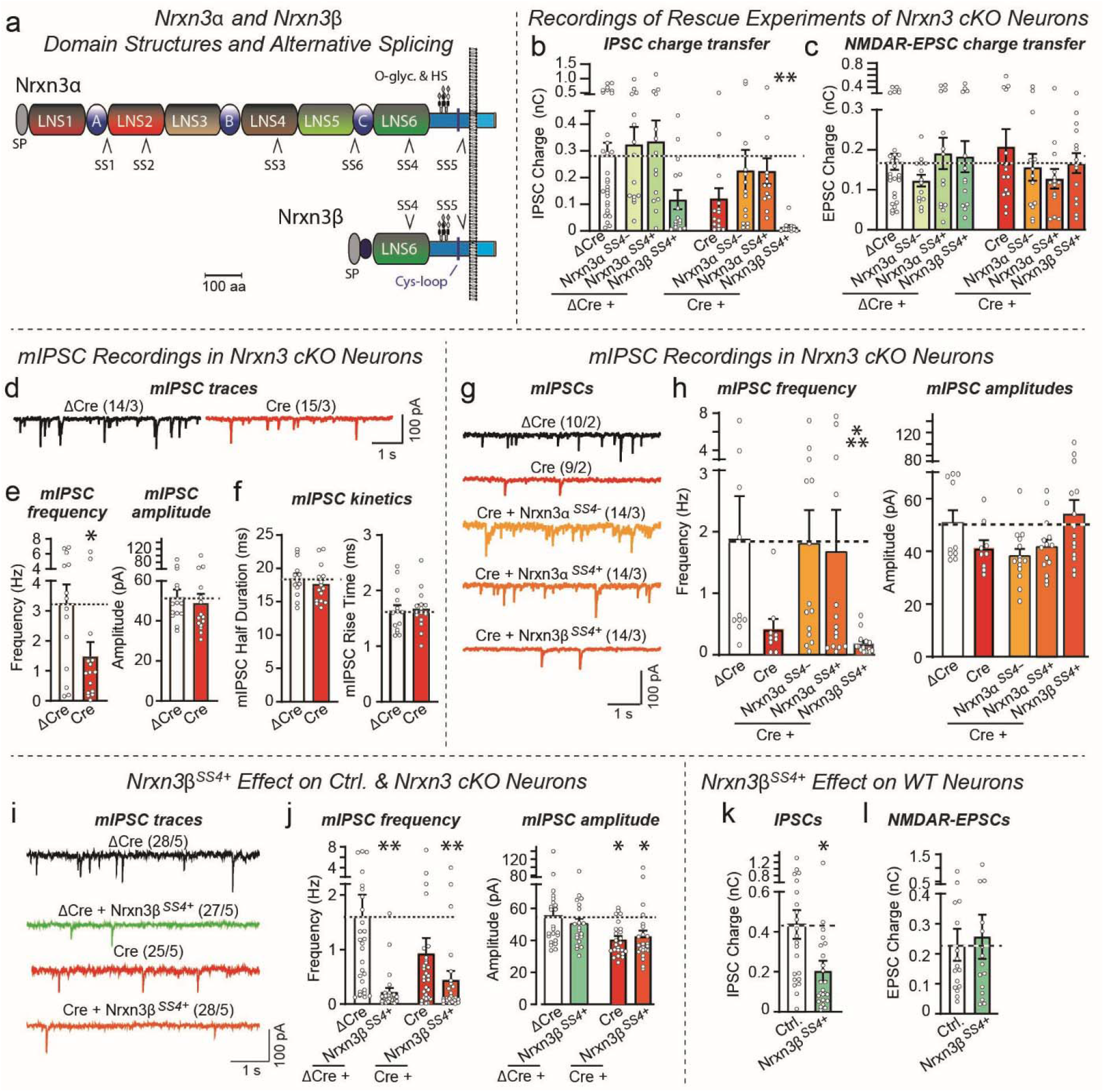
Further analyses of Nrxn3α and Nrxn3β and their SS4-splice variants confirm that both Nrxn3α-SS4+ and Nrxn3α-SS4-rescue impaired GC→MC inhibitory synaptic transmission in *Nrxn3*-deficient cultured OB neurons, whereas Nrxn3β suppresses, instead of rescuing, synaptic transmission at these synapses, and inhibits GC→MC synapses even in control. **a**. Schematic of Nrxn3α and Nrxn3β domain structures and their sites of alternative splicing (SP, signal peptide; SS1-SS6, alternatively spliced sequences #1-6; A, B, C, first, second, and third EGF-like domain; O-glyc. & HS, O-glycosylation and heparan-sulfate attachment sequences). **b** & **c**. Quantification of the synaptic charge transfer during evoked IPSCs confirms that the conditional *Nrxn3* deletion severely impairs evoked IPSCs in a manner that can be rescued by Nrxn3α but not Nrxn3β (b), but that the conditional deletion of *Nrxn3* has no effect on evoked NMDAR-EPSCs (c). Data are from the same experiments as shown in Figure 1a, 1b, 1e & 1f. **d**-**f**. Conditional deletion of *Nrxn3* in cultured OB neurons decreases the frequency of spontaneous mIPSCs by ∼60% without significantly altering the amplitude or kinetics of mIPSCs (d, sample traces; e, summary graphs of the mIPSC frequency and amplitudes; f, summary graphs of the mIPSC decay and rise times). **g** & **h**. Conditional deletion of *Nrxn3* in cultured OB neurons massively decreases spontaneous mIPSCs; this decrease is rescued by Nrxn3α but not Nrxn3β (g, sample traces; h, summary graphs of mIPSC frequency and amplitudes). **i** & **j**. Independent experiments documenting that expression of Nrxn3β-SS4+ with an insert at SS4 suppresses the mIPSC frequency in both wild-type and *Nrxn3*-deficient cultured OB neurons (i, sample traces; j, summary graphs of the mIPSC frequency and amplitudes). Note that the small but statistically significant decrease in mIPSC amplitudes observed in this experiment after the *Nrxn3* deletion is not observed in the experiments of panels e and h of this figure, although there was a trend towards a decrease in h. **k** & **l**. Quantification of the synaptic charge transfer during evoked IPSCs confirms that the expression of Nrxn3β-SS4+ in wild-type neurons severely impairs evoked IPSCs in a dominant-negative fashion (k) but has no effect on evoked NMDAR-EPSCs (l). Data are from the same experiments as shown in Figure 1c, 1d, 1g & 1h. Numerical data are means ± SEM; n’s (cells/experiments) are indicated above the sample traces and apply to all graphs in an experimental series. Statistical analyses were performed with a one-way analysis of variance (ANOVA) with Dunnett’s multiple comparison test (b, c, h, and j) or a Welch’s *t* test (e, f, k, and l), with * = p<0.05, ** = p<0.01, and *** = p<0.001.

**Figure S2:**
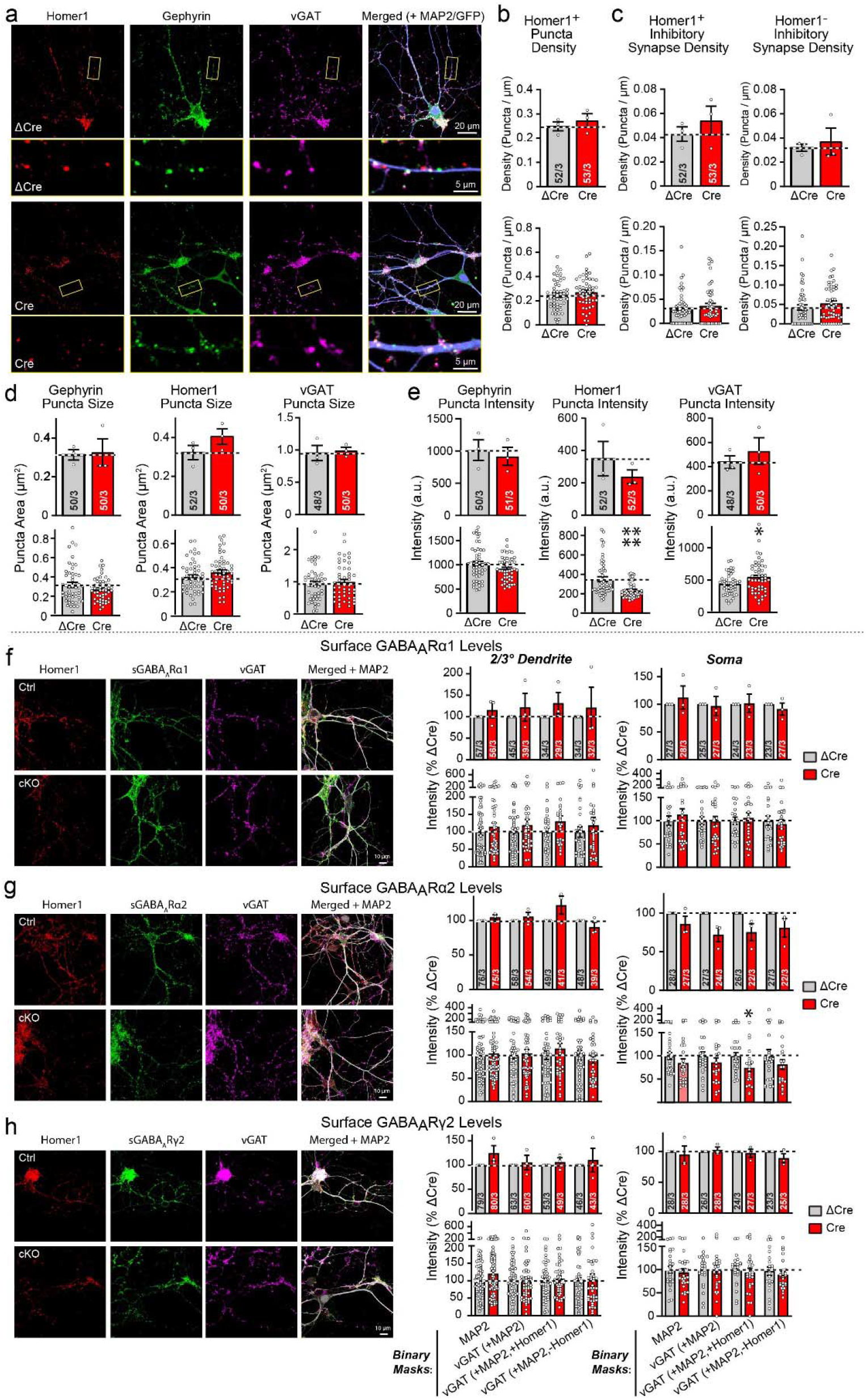
The *Nrxn3* deletion does not alter the density of GC→MC synapses in cultured OB neurons (a-e), nor does it alter the surface levels of postsynaptic GABA_A_-receptors (f-h) **a.** Further representative images related to the data of Fig. 1i & 1j. **b & c.** Additional quantifications of immunocytochemical analyses of cultured OB neurons in which larger mitral/tufted cells form dendrodendritic synapses with smaller granule cells and other interneurons, confirming that the synapse density is not altered by deletion of *Nrxn3*. Summary graphs show quantifications of the density of Homer1+ excitatory synapses (b, Homer1-positive), putative reciprocal synapses (c, synapses that are both excitatory and inhibitory as evidenced by Homer1+, gephyrin+, and vGAT+ signals), and exclusively inhibitory synapses (c, gephyrin+ & vGAT+ but Homer1-). The logic of these quantifications is that reciprocal synapses composed of adjacent excitatory and inhibitory synaptic junctions in opposite orientations contain both Homer1 and gephyrin as excitatory and inhibitory synaptic markers, whereas pure excitatory or pure inhibitory synapses contain only one or the other marker. Data are shown as averaged per experiment (top) or per ROI (bottom) since the latter is the standard of the field but can create artifactual statistical significance since the statistics is not based on the number of experiments, and a single experiment can become highly statistically significant. Note that there is no meaningful difference between control and *Nrxn3*-deficient neurons in any of the quantified parameters. **d & e.** The *Nrxn3* deletion has no effect on the size (d) or intensity (e) of synaptic puncta (left, gephyrin; middle, Homer1; right, vGAT), suggesting that it doesn’t cause a major structural reorganization of synapses. Data are shown as averaged per independent culture (top) or per ROI (bottom). **f-h**. The surface expression of several GABA_A_R subunits, namely GABA_A_Rα1 (f), GABA_A_Rα2 (g) and GABA_A_Rγ2 (h) is not altered by the *Nrxn3* deletion. For each set of data, representative images are shown on the left and summary graphs on the right. Receptor levels were measured by staining non-permeabilized neurons with antibodies to the indicated receptors, and were quantified at secondary and tertiary dendrites (left summary graphs) and over the soma (right summary graphs). Receptors levels monitored as background-subtracted fluorescence intensity are quantified in areas defined by binary masks of other stains including the dendrite and various combinations of synaptic markers. Data are shown as averaged values per experiment (top) or per ROI (bottom) for the reasons explained in the legend to panels b & c. Numerical data are means ± SEM; n’s (images/experiments) are indicated in the summary graph bars and apply to all graphs in an experimental series. Statistical analyses were performed using a Welch’s *t* test, with * = p<0.05, ** = p<0.01, *** = p<0.001. **** = p<0.0001.

**Figure S3:**
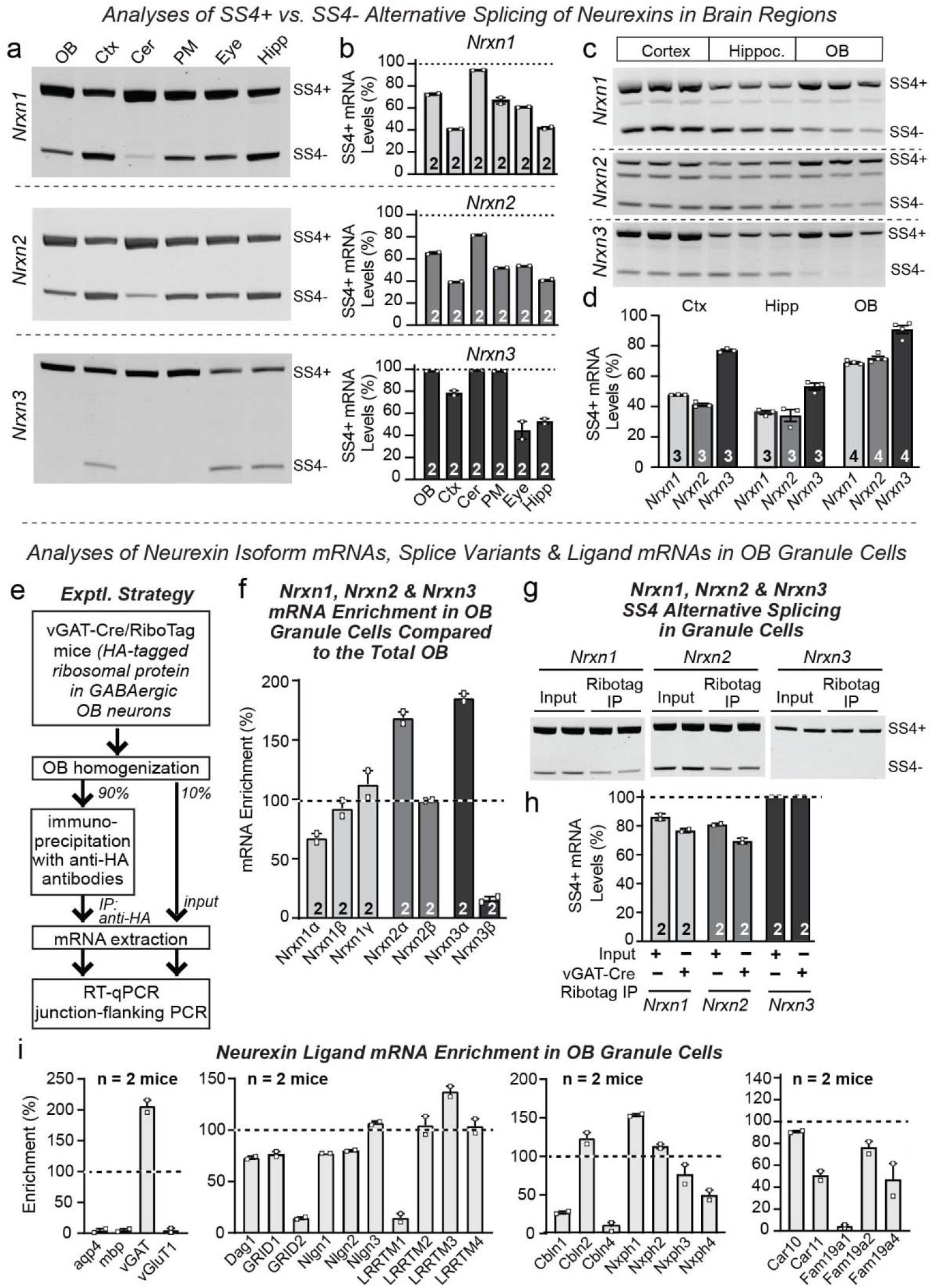
RT-PCR analyses of the alternative splicing of neurexins at SS4 in different brain regions (a-d), of the mRNA levels of neurexins and of their SS4 alternative splicing in OB granule cells (e-h), and of the mRNA levels of selected cis- and trans-ligand of neurexins in OB granule cells (i) reveals a regulated expression pattern of neurexin mRNAs and their SS4 splice variants. **a & b**. *Nrxn1*-*Nrxn3* mRNAs exhibit brain region-specific differences in SS4 alternative splicing inclusion as determined by junction-spanning RT-PCR of RNA isolated from the indicated brain regions, followed by electrophoretic analyses and quantification of the PCR products (a, representative gels; b, summary graphs of the SS4+ RNA as percent of the total). Abbreviations used: OB, olfactory bulb; Ctx, cortex; PM, pons and medulla; Cer, cerebellum; Hipp, hippocampus. **c & d**. *Nrxn1*-*Nrxn3* mRNAs also exhibit brain region-specific differences in SS4 alternative splicing when analyzed by junction-spanning RT-PCR of RNA isolated from dissociated cultures from the cortex (Cx), hippocampus (Hipp), and olfactory bulb (OB) of mice (c, representative images; d, summary graphs). **e.** Experimental strategy to isolate translating mRNAs from OB granule cells using RiboTag mice and for analysis of these mRNAs. The strategy is based on the fact that the vast majority of inhibitory neurons in the OB (>90%) are granule cells. As a result, RiboTag purification of mRNAs from vGAT-positive neurons primarily isolates granule cell mRNAs. **f**. Nrxn2α and Nrxn3α mRNAs are enriched in OB granule cells compared to total RNA of the OB, whereas Nrxn3β mRNAs are de-enriched, and Nrxn1γ and Nrxn2β mRNAs are neither enriched nor de-enriched. Data show relative abundance of indicated mRNAs in OB granule cells normalized to the total mRNAs, and do not depict absolute mRNA levels. **g** & **h**. Most (70-80%) Nrxn1α, Nrxn1β, Nrxn1α, and Nrxn1β mRNAs, but nearly all (∼100%) of Nrxn3α and Nrxn3β mRNAs, are expressed as SS4+ variants in OB granule cells (g, representative image of junction-flanking PCR analysis of total OB mRNA (input) and vGAT-Cre/Ribotag-isolated OB granule cell mRNA [immunoprecipitation (IP): anti-HA]; left, summary graph]; h, quantification of the relative abundance of SS4+ mRNAs). **i.** Validation of the purification of mRNAs from OB granule cells by quantifications of the enrichment of markers for specific OB cell populations (left summary graph; aqp, aquaporin as an astrocyte marker; mbp, myelin basic protein as an oligodendrocyte marker; vGluT1 and vGAT as excitatory and inhibitory neuron markers, respectively), and measurements of the enrichment of selected neurexin ligand mRNAs in OB granule cells (middle and right summary graphs). Numerical data are means ± SEM; n’s (cells/experiments) are indicated in the summary graph bars and apply to all graphs in an experimental series.

**Figure S4:**
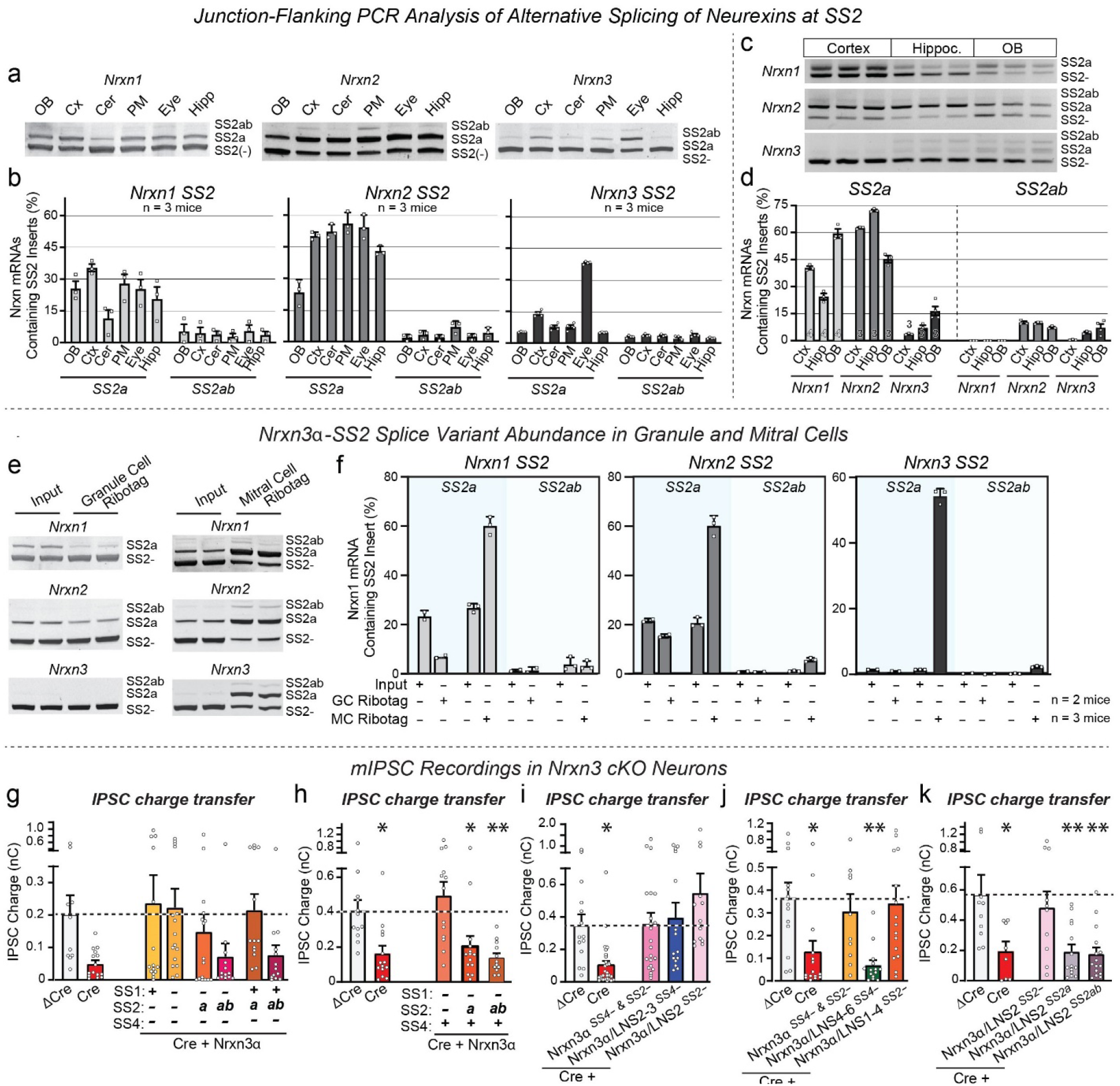
RT-PCR analyses of the alternative splicing of *Nrxn1-Nrxn3* mRNAs at SS2 in different brain regions (a-d) and in OB mitral/tufted and granule cells (e & f), and confirmation of the hierarchical SS2 and SS4 splice code of *Nrxn3* function in inhibitory OB synapses by analyses of the synaptic charge transfer in rescue experiments (g-k) **a** & **b**. Analyses of the alternative splicing of *Nrxn1*-*Nrxn3* mRNAs in various brain regions at SS2, which is expressed in three variants: Two more common variants lacking (SS2-) or containing an 8 residue insert (SS2a), and a very rare variant that includes not only the 8-residue ‘a’ insert, but also an additional 7-residue ‘b’ insert (SS2ab; Ullrich et al., 1995). Splice variants were measured in mRNAs from the indicated brain regions via junction-spanning RT-PCR followed by quantification of the splicing patterns on gels. As shown, a sizable fraction of *Nrxn1* mRNAs (∼15-30%) and of *Nrxn2* mRNAs (∼50%) contain an ‘a’ insert in SS2 in most brain regions, but only a small fraction of *Nrxn3* mRNAs (<15%) contains such an insert with the exception of the eye (∼40%). In all brain regions, mRNAs with ‘ab’ inserts are exceedingly rare (a, representative gels of junction-spanning RT-PCR products; b, summary graphs of the percentages of the SS2a and SS2ab mRNAs of the indicated neurexins). **c** & **d**. Analyses of neurexin-SS2 alternative splicing by junction-spanning RT-PCR in dissociated cultures from cortex (Ctx), hippocampus (Hipp), and OB confirms that most *Nrxn3* mRNAs lack an insert in SS2, with ∼30% of *Nrxn1* mRNAs, ∼50% of Nrxn2 mRNAs, but <20% of *Nrxn3* mRNAs expressed as SS2+ mRNAs (c, representative gels; d, summary graphs of the levels SS2 variants, SS2a and SS2ab, expressed as a % of total transcripts). **e** & **f**. Junction-spanning RT-PCR measurements of SS2-, SS2a, and SS2ab splice variants in mRNAs isolated by RiboTag IPs from OB mitral/tufted and granule cells uncovers a distinctive pattern of cell type-specific alternative splicing whereby granule cells selectively express SS2-neurexin variants, whereas mitral/tufted cells express SS2+ variants (e, representative splice-junction PCR gels; f, summary graphs of the percentages of SS2a and SS2ab mRNA). Note that SS2+ mRNAs in mitral/tufted cells could serve to inhibit cis-interactions with mitral cell dystroglycan. Cell-specific mRNAs were purified and their purity validated as described in Fig. S3e and Wang et al., 2021, with granule cell mRNA isolated from RiboTag mice carrying vGAT-Cre and mitral/tufted cell mRNA isolated from RiboTag mice carrying TBet-Cre. **g** - **k**. Quantifications of the synaptic charge transfer during evoked IPSCs confirm that minimal Nrxn3α-LNS2 constructs lacking an insert in SS2 fully rescue the suppression of evoked IPSCs induced by the conditional deletion of *Nrxn3* in cultured OB neurons. Experiments are the same as those shown in Figure 1k-1m and Figure 2b-2g. Numerical data are means ± SEM; n’s (images/experiments) are indicated in the summary graph bars (a-f) and apply to all graphs in an experimental series, or are described in the main figures. Statistical analyses were performed with a one-way analysis of variance (ANOVA) with Dunnett’s multiple comparison test (g-k), with * = p<0.05, ** = p<0.01, and *** = p<0.001.

**Figure S5:**
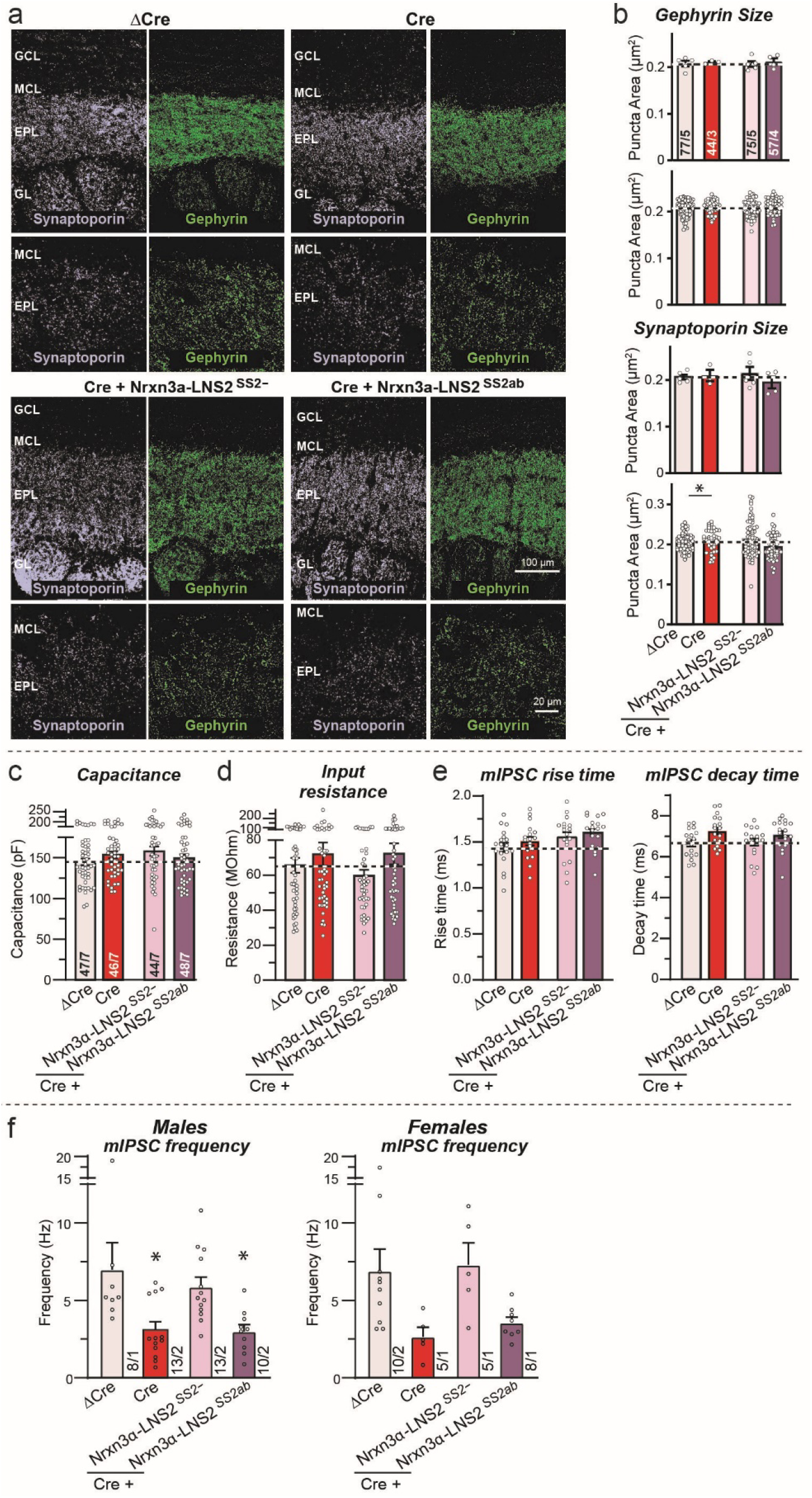
Further data confirming that the Nrxn3 deletion in the OB in vivo does not lower synapse numbers (a & b), but that the severe impairment in inhibitory GC→MC synapses induced by the Nrxn3 deletion in vivo can be rescued with the Nrxn3α^LNS2-^, but not the Nrxn3α^LNS2ab^, construct. **a** & **b**. Conditional deletion of *Nrxn3* and expression of Nrxn3α^LNS2-^ and Nrxn3α^LNS2ab^ rescue constructs in the OB in vivo do not alter the size of reciprocal mitral/granule cell synapses as analyzed by quantitative immunohistochemistry for a presynaptic (synaptoporin) and postsynaptic marker (gephyrin) (a, sample images; b, summary graphs of puncta size). Note that puncta sizes in b are plotted both as per animal (top) and as per region-of-interest (bottom) because the latter approach is the standard of the field but artifactually boosts statistical significance independent of the actual number of experiments. Data complement those shown in Fig. 3. **c** & **d**. Measurements of the cell capacitance (c) and input resistance (d) show that the conditional deletion of *Nrxn3* and the expression of Nrxn3α^LNS2-^ and Nrxn3α^LNS2ab^ rescue constructs does not significantly affect the basic electrical properties of neurons. Data complement those shown in Fig. 4. **e**. Quantifications of the mIPSC rise and decay times uncover no changes induced by the conditional deletion of *Nrxn3* or the expression of Nrxn3α^LNS2-^ and Nrxn3α^LNS2ab^ rescue constructs in mitral cells in vivo. Data complement those shown in Fig. 4a-c. **f**. Plots of the data shown in Figure 4a-4b separately for male and female mice to demonstrate that both genders exhibit similar phenotypes. Numerical data are means ± SEM; n’s (images/experiments) are indicated in the summary graph bars and apply to all graphs in an experimental series. Statistical analyses were performed with a one-way analysis of variance (ANOVA) with Dunnett’s multiple comparison test comparing all samples to the ΔCre control condition, with * = p<0.05 and ** = p<0.01.

**Figure S6:**
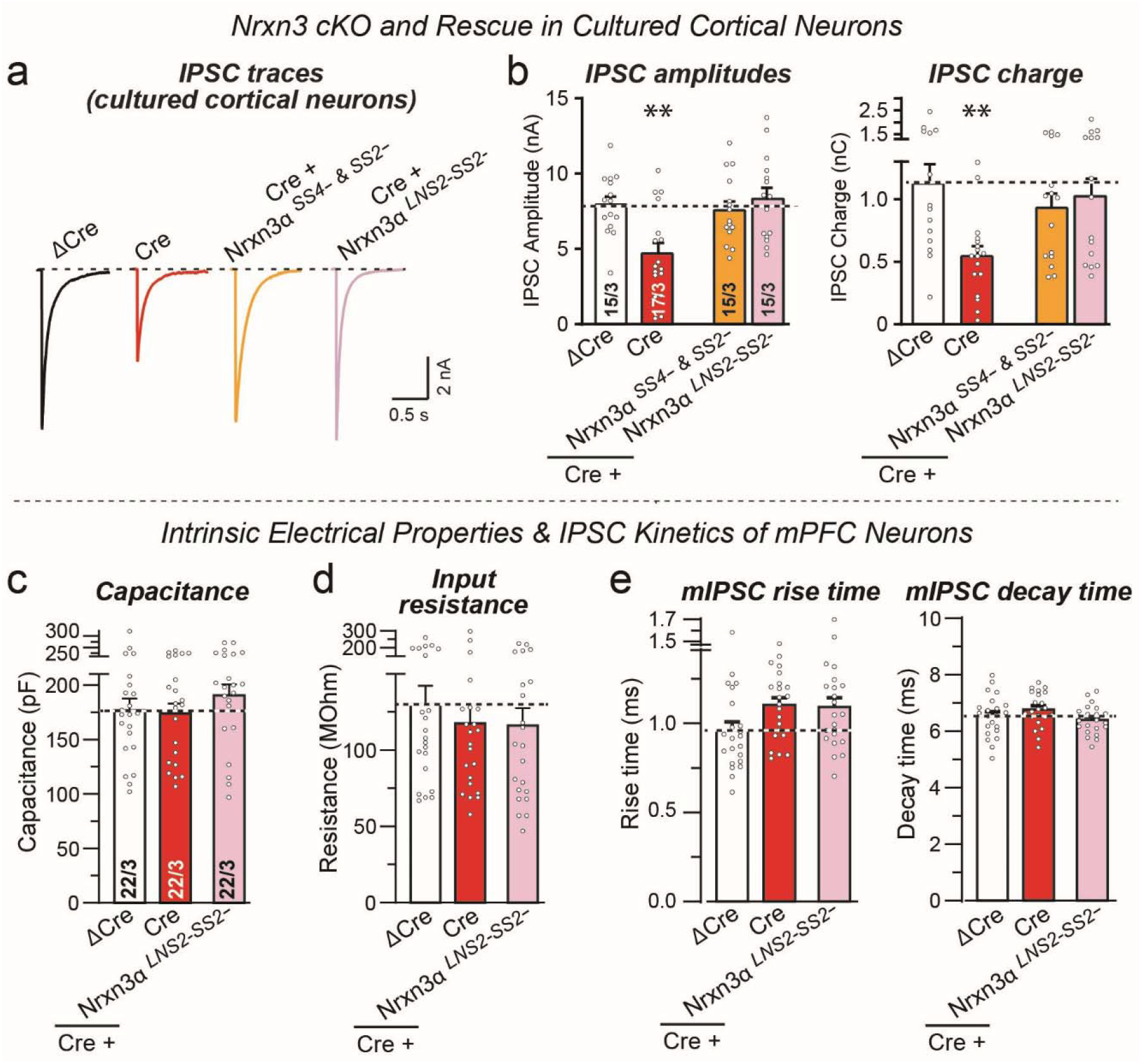
Conditional *Nrxn3* deletion of in cultured cortical neurons impairs evoked inhibitory synaptic transmission (a & b), and conditional *Nrxn3* deletion in the mPFC in vivo does not detectably alter the passive electrical properties of pyramidal neurons (c & d) or the kinetics of mIPSCs (e) **a** & **b**. Conditional deletion of *Nrxn3* in cultured cortical mouse neurons significantly decreases the amplitude and charge transfer of evoked IPSCs. Neurons were cultured from newborn *Nrxn3* cKO mice, infected with lentiviruses expressing ΔCre (control) or Cre at 4 days in culture, and analyzed at 14-16 days in culture (a, representative traces; b, summary graphs of the IPSC amplitude and charge transfer). The experiments provided the rationale for performing in vivo experiments shown in Fig. 5. **c** & **d**. Measurements of the cell capacitance (c) and input resistance (d) show that the conditional deletion of *Nrxn3* and the expression of Nrxn3α^LNS2-^ and Nrxn3α^LNS2ab^ rescue constructs in the mPFC in vivo does not significantly affect the basic electrical properties of neurons. Data complement those shown in Fig. 5. **e**. Quantifications of the mIPSC rise and decay times uncover no changes induced by the conditional deletion of *Nrxn3* or the expression of Nrxn3α^LNS2-^ and Nrxn3α^LNS2ab^ rescue constructs in the mPFC in vivo. Data complement those shown in Fig. 5. Numerical data are means ± SEM; n’s (cells/experiments) are indicated in the summary graph bars and apply to all graphs in an experimental series. Statistical analyses were performed with a one-way analysis of variance (ANOVA) with Dunnett’s multiple comparison test comparing all samples to the ΔCre control condition, with * = p<0.05, ** = p<0.01, and *** = p<0.001.

**Figure S7:**
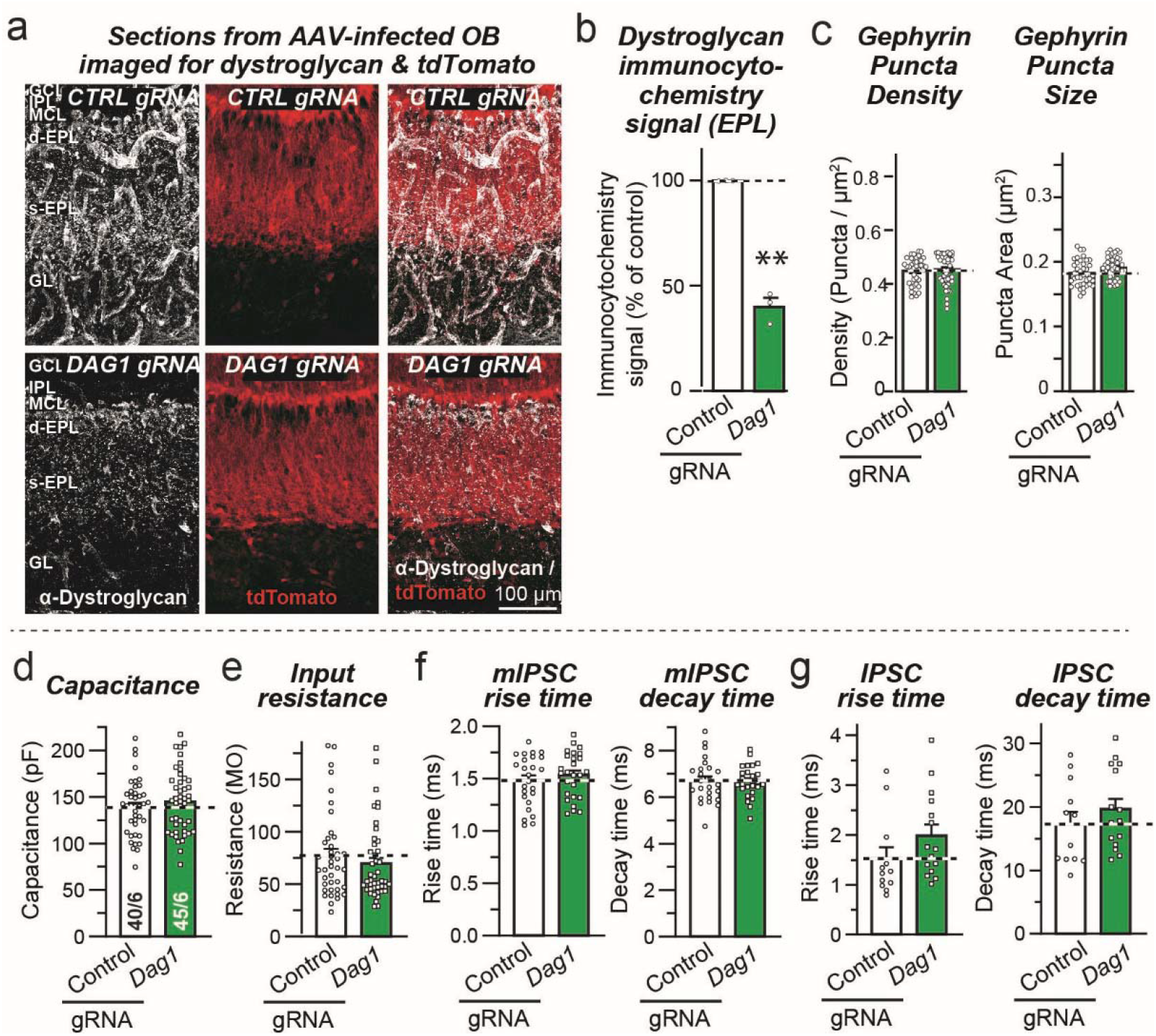
CRISPR-mediated in vivo deletion of dystroglycan (*Dag1*) in the OB is efficient (a, b), doesn’t change synapse numbers (c), and has no major effects on the intrinsic electrical properties (d, e) or the mIPSC (f) or evoked IPSC kinetics (g) monitored in mitral cells in acute slices. **a** & **b**. Analyses of the effect of the CRISPR-mediated deletion of dystroglycan (*Dag1*) in the OB by immunocytochemistry, demonstrating that the CRISPR manipulation suppresses expression of ∼60% of dystroglycan in the external plexiform layer (EPL) (a, representative images of OB sections stained for dystroglycan and tdTomato (to mark AAV-infected cells, see Fig. 7a-d); b, summary graph of the dystroglycan staining intensity demonstrating loss of dystroglycan). **c**. The dystroglycan (*Dag1*) deletion has no effect on the density or size of gephyrin+ synaptic puncta. Summary graphs depict an analysis of gephyrin+ synaptic puncta in the OB as a function of the dystroglycan deletion quantified per region-of-interest instead of per experiment to demonstrate that even with pseudo-replicates, the dystroglycan deletion does not alter synapse density. Experiment is the same as shown in Figure 7c & 7c. **d** & **e**. Summary graphs of the capacitance (d) and input resistance (e) of mitral cells as a function of the dystroglycan deletion in the OB in vivo described in Figure 7f-7n demonstrate that the dystroglycan deletion does not detectably alter the size and membrane properties of the neurons. **f**. Summary graphs of the mIPSC rise and decay times of mitral cells as a function of the dystroglycan deletion in the OB in vivo described in Figure 7f-7h demonstrate that the dystroglycan deletion does not cause a detectable change in the mIPSC kinetics. **g**. Summary graphs of the IPSC rise and decay times monitored in mitral cells as a function of the in vivo dystroglycan deletion in the OB described in Figure 7i-7n demonstrate that the dystroglycan deletion does not detectably impair the IPSC kinetics. Numerical data are means ± SEM; n’s (cells or images/experiments) are indicated in the summary graph bars and apply to all graphs in an experimental series. Statistical analyses were performed using Student’s t-test, with * = p<0.05, ** = p<0.01, and *** = p<0.001.

**Figure S8:**
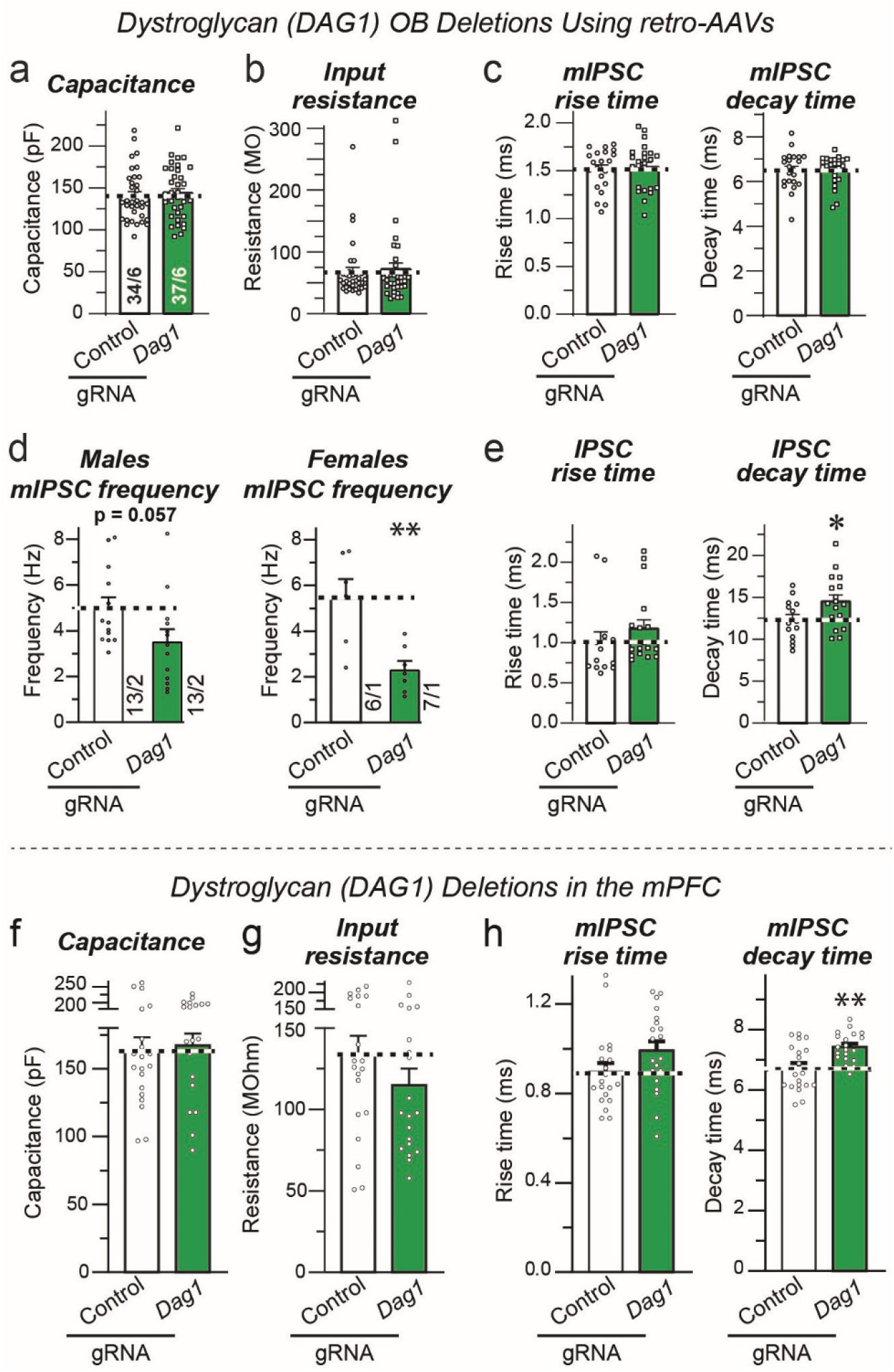
Further characterization of the CRISPR-mediated in vivo deletion of dystroglycan (*Dag1*) in mitral cells on the morphological and electrophysiological properties of GC→MC synapses, as obtained by retrograde expression of gRNAs in mitral cells via infection of the piriform cortex with retro-AAVs that produces a deletion of dystroglycan only in mitral/tufted cells projecting to the piriform cortex (a-e), and of the in vivo deletion of dystroglycan in the mPFC (f-h) **a** & **b**. Summary graphs of the capacitance (a) and input resistance (b) of mitral cells as a function of the dystroglycan deletion specifically only in mitral cells using retro-AAVs injected into the piriform cortex as described in Figure 8a demonstrate that the dystroglycan deletion does not detectably cause a major change in the size and membrane properties of the neurons. Data correspond to the recordings shown in Figure 8f-8n. **c.** Summary graphs of the mIPSC rise and decay times of mitral cells as a function of the dystroglycan deletion in the OB in vivo described in Figure 8f-8h demonstrate that the dystroglycan deletion does not induce a detectable change in the mIPSC kinetics. **d.** Plots of the data shown in Figure 8f-8h separately for male and female mice to demonstrate that both sexes exhibit similar phenotypes. **e.** Summary graphs of the IPSC rise and decay times of mitral cells as a function of the dystroglycan deletion in the OB in vivo described in Figure 8i-8n demonstrate that the dystroglycan deletion does not give rise to a detectable change in the IPSC kinetics. **f** & **g**. Summary graphs of the capacitance (f) and input resistance (g) of pyramidal neurons in the mPFC as a function of the dystroglycan deletion in the mPFC in vivo described in Figure 8o-8r demonstrate that the dystroglycan deletion does not cause a major change in the size and membrane properties of the neurons. **h**. Summary graphs of the mIPSC rise and decay times of layer 5 pyramidal neurons as a function of the dystroglycan deletion in the mPFC in vivo described in Figure 8p-8r demonstrate that the dystroglycan deletion does not detectably cause a major change in the mIPSC kinetics. Numerical data are means ± SEM; n’s (cells or images/experiments) are indicated in the summary graph bars and apply to all graphs in an experimental series. Statistical analyses were performed using Student’s t-test, with * = p<0.05, ** = p<0.01, and *** = p<0.001.

